# RIG-I uses its intrinsically disordered CARDs-Helicase linker in RNA proofreading functions

**DOI:** 10.1101/2021.09.23.461459

**Authors:** Brandon D. Schweibenz, Swapnil C. Devarkar, Mihai Solotchi, Jie Zhang, Bruce D. Pascal, Patrick R. Griffin, Smita S. Patel

## Abstract

The innate immune receptor RIG-I provides the first line of defense against viral infections. Viral RNAs are recognized by the C-terminal domain (CTD) of RIG-I, but the RNA must engage the helicase domain to release the signaling domain CARDs from their autoinhibitory CARD2:Hel2i interactions. Because the helicase lacks RNA specificity, there must be mechanisms to proofread RNAs entering the helicase domain. Although such mechanisms are crucial in preventing aberrant immune responses by non-specific RNAs, they remain largely unknown. This study reveals a previously unknown proofreading mechanism that RIG-I uses to ensure the helicase engages RNAs chosen explicitly by the CTD. A crucial part of this mechanism involves the intrinsically disordered CARDs-Helicase Linker (CHL), which uses its negatively charged regions to electrostatically antagonize incoming RNAs. In addition to RNA gating, CHL is essential in stabilizing the CARD2:Hel2i interface. The CHL and CARD2:Hel2i interface work together, establishing a tunable gating mechanism that allows CTD-chosen RNAs to bind into the helicase while blocking non-specific RNAs. With its critical regulatory functions, CHL represents a novel target for RIG-I-based therapeutics.

## Introduction

RIG-I (Retinoic Acid Inducible Gene-I) is an innate immune receptor responsible for surveilling the cytoplasm for viral RNAs (Yoneyama *et al*, 2005). RIG-I recognizes short blunt-ended double-stranded (ds) RNAs with 5’-triphosphate (5’ppp), 5’-diphosphate (5’pp), and 5’-m7G cap as PAMPs (pathogen-associated molecular pattern) (Devarkar *et al*, 2016; Goubau *et al*, 2014; Jiang *et al*, 2011; Schuberth-Wagner *et al*, 2015). Such PAMP features are not present in endogenous RNAs but found in many RNA genomes and most replication intermediates of negative-strand and positive-strand RNA viruses (Devarkar *et al*., 2016; Hu *et al*, 2017; Kato *et al*, 2006; Rehwinkel *et al*, 2010; Schuberth-Wagner *et al*., 2015; Stumper *et al*, 2005). Upon recognizing viral RNAs, RIG-I initiates a signaling cascade that culminates in Type I interferon response, efficiently controlling viral infections (Poeck *et al*, 2010; Stumper *et al*., 2005; Yoneyama *et al*, 2004).

Structural studies show that RIG-I has flexibly-linked domains that can switch between active and inactive states to respond appropriately to viral RNAs (Figure 1A) (Civril *et al*, 2011; Devarkar *et al*., 2016; Jiang *et al*., 2011; Kowalinski *et al*, 2011; Luo *et al*, 2011; Luo *et al*, 2012). The core helicase domain contains two helicase subdomains, Hel1 and Hel2, and a nested Hel2i, flanked by a tandem array of signaling domain CARDs (Caspase Activation and Recruitment Domains) and a PAMP recognition C-terminal domain (CTD). When RIG-I is not bound to RNA, the helicase domain is in an open conformation, inactive in ATP hydrolysis. In this conformation, the CARDs are sequestered by Hel2i through CARD2:Hel2i interactions and inactive in signaling (**Figure 1A**). When RIG-I fully engages the RNA, the induced conformational change activates the ATPase and disrupts the CARD2:Hel2i interface to release the CARDs from their autoinhibitory interactions (Dickey *et al*, 2019; Zheng *et al*, 2018; Zheng *et al*, 2015). The exposed CARDs can initiate the immune response by interacting with downstream adapter proteins (Peisley *et al*, 2014; Wu *et al*, 2014).

**Figure 1.**
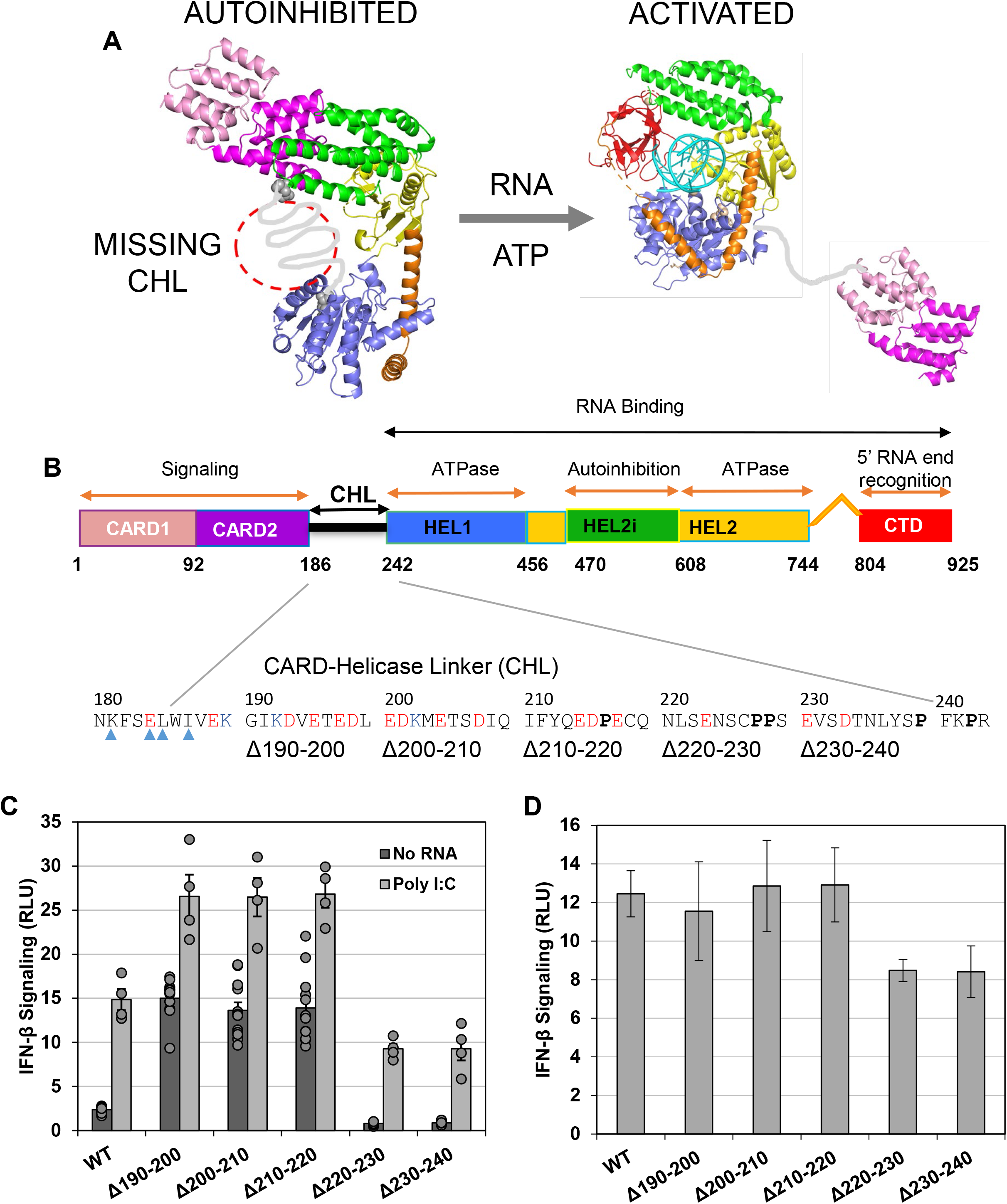
RIG-I CARDs-Helicase Linker (CHL) regulates RIG-I signaling. (A) In the autoinhibited state of RIG-I, the helicase is an open conformation and the CARDs are interacting with Hel2i. The missing CHL in gray passes near the RNA binding groove of the helicase domain as its spans the 40 Å distance between the C-terminus of CARD2 and N-terminus of Hel1. PAMP RNA and ATP binding induces a conformational change to close the helicase subdomains around the ligands and release the CHL and CARDs. Models were generated from crystal structures (PDB ID: 4A2W, left; composite model of two crystal structures PDB IDs: 4NQK and 5E3H, right and colored as in (B). (B) Domain schematic of RIG-I with the CHL sequence highlighted. Each CHL deletion mutant is indicated below the sequence. Blue arrows point out the CARD2 residues known to contact the Hel2i domain to form the CARD2:Hel2i interface in the autoinhibited duck RIG-I. Negative and positive charged amino acids are labeled red and blue, respectively. Proline resides are shown in bold. (C) Cell signaling experiments testing the contribution of CHL to RIG-I activity. Either no RNA or poly I:C RNA (a known RIG-I stimulator) was transfected. Circles indicate individual replicates (six for No RNA, four for poly I:C), and the error bar is SEM. (D) Cell signaling experiments from (C), except the No RNA background signal has been subtracted from the poly I:C signal for each construct to determine the contribution of poly I:C to RIG-I signaling activity.

It is crucial that viral RNAs and not self RNAs, abundantly present in the cytoplasm, activate RIG-I. Aberrant immune responses initiated by RIG-I are harmful and lead to autoinflammatory disorders (Crowl *et al*, 2017; Roers *et al*, 2016). RIG-I has evolved several mechanisms to discriminate self and non-self, including its ATPase activity to power its translocation and dissociation from non-PAMP RNAs (Devarkar *et al*, 2018). PAMP RNAs are recognized specifically by the CTD of RIG-I, which provides the first layer of RNA proofreading (Cui *et al*, 2008; Lu *et al*, 2010; Ramanathan *et al*, 2016; Vela *et al*, 2012; Wang *et al*, 2010). The CTD binds PAMP RNA ends in RIG-I’s autoinhibited state with CARDs still engaged with the Hel2i (Devarkar *et al*., 2018). To activate the signaling CARDs, the chosen RNA must be loaded into the helicase domain. Unlike CTD, the helicase domain lacks RNA specificity and hence can bind non-specific RNAs. How RIG-I ensures that the helicase domain binds RNAs selected by the CTD has remained unclear. Previous studies suggested that the CARD2:Hel2i interface regulates RNA binding into the helicase domain, but the underlying mechanism was not known (Ramanathan *et al*., 2016; Vela *et al*., 2012). We discovery serendipitously that RIG-I uses the ~56 amino acids flexible linker, connecting the CARDs to the helicase domain, as this crucial regulatory region to control the helicase domain activities of RNA binding and CARDs activation. The linker has been considered to be a passive region, thus far.

The 190-240 polypeptide constituting the CARDs-Helicase Linker (CHL) is predicted to be intrinsically disordered (Oates *et al*, 2013). Consistent with its flexibility, CHL was not resolved in the autoinhibited duck RIG-I structure (Kowalinski *et al*., 2011). However, the structure suggests that the missing CHL is located close to the helicase domain (**Figure 1A**). We found that small deletions in CHL resulted in hyperactive RIG-I signaling responses, indicating CHL’s role in regulating CARDs exposure. We created several CHL region mutants and characterized them using a wide range of biochemical and biophysical methods, including hydrogen-deuterium exchange and mass spectrometry (HDX-MS), equilibrium RNA binding, and stopped-flow kinetics to understand the regulatory role of CHL. These studies revealed two essential functions of the CHL. Firstly, CHL stabilizes the CARD2:Hel2i interface to keep RIG-I in an autoinhibited state in the absence of PAMP RNA. Secondly, CHL electrostatically gates the helicase domain to proofread RNAs entering the helicase domain. Both of these functions depend on the negatively charged regions in the CHL. We show how CHL and CARD2:Hel2i interface work at different steps in the RNA binding pathway to block non-specific RNAs while allowing RNAs chosen by the CTD to form a stable complex. The newly discovered role of the CHL can be leveraged in RIG-I based therapeutics.

## RESULTS

### CHL deletions cause hyperactive signaling response in the absence of PAMP RNA

To investigate the role of CHL in RIG-I signaling, we made segmental deletions of 11 amino acids in the CHL region between 190-240 (**Figure 1B**). The expression and signaling activity of CHL deletion mutants and wild-type (WT) RIG-I was tested in HEK-293T cells using the dual-luciferase IFN-β promoter reporter assay in the absence and presence of PAMP RNA. The signaling activity of WT RIG-I without PAMP RNA was minimal, indicative of tightly regulated CARDs (**Figure 1B and Figure EV1**). To our surprise, three of the five CHL deletion mutants showed a hyperactive signaling response even in PAMP RNA’s absence. The deletions closest to the CARDs, Δ190-200, Δ200-210, and Δ210-220, showed ~10 fold higher signaling responses than WT RIG-I (**Figure 1C**). The deletion mutants closest to Hel1, Δ220-230 and Δ230-240, behaved like WT RIG-I. Interestingly, CHL deletions did not impair the signaling response in the presence of PAMP RNA. We observed a normal PAMP-stimulated signaling response from CHL mutants with Poly I:C (**Figures 1C and D**). The RIG-I mutants, Δ190-200, Δ200-210, and Δ210-220, were activated to a similar extent as WT RIG-I. The Δ220-230 and Δ230-240 mutants were activated to a slightly lower level.

These results provided the first evidence that CHL is a regulatory region that suppresses the aberrant signaling activity of RIG-I in the absence of PAMP RNA.

### CHL stabilizes the CARD2:Hel2i interface to minimize CARDs exposure and signaling responses from non-PAMP RNA

There are two possible ways the CHL deletion could activate RIG-I signaling in the absence of PAMP RNA. One, CHL deletions destabilize the CARD2:Hel2i interface to expose the signaling CARDs spontaneously. Two, CHL deletions impair RIG-I’s RNA discrimination mechanism, enabling self-RNAs to bind and activate the signaling domain CARDs.

To test the first possibility, we used differential HDX-MS analysis, a powerful technique used to monitor CARDs exposure in RIG-I previously (Zheng *et al*, 2019; Zheng *et al*., 2018; Zheng *et al*., 2015). Differential HDX-MS analysis of WT and Δ190-200 RIG-I in the absence of PAMP RNA showed increased deuterium exchange in many regions of RIG-I due to the CHL deletion (**Figures 2A and 2B, Figure EV2**). A higher degree of deuterium exchange is indicative of greater peptide backbone flexibility. The Δ190-200 RIG-I showed increased peptide backbone flexibility in the CARDs (aa 1-186), Hel1 (aa 410-460), and several areas of Hel2 (aa 695-756, 785-803) and Hel2i (aa 566-574 and 522-539), indicating a global effect of CHL deletion that destabilizes the autoinhibited conformation of RIG-I. Interestingly, increased deuterium exchange was observed at the CARD2:Hel2i interface, comprising the CARD2 latch peptide (aa 103-114) and the interacting Hel2i regions (aa 566-574 and 522-539). These results provide strong evidence for the first possibility that the 190-200 CHL deletion disrupts the CARD2:Hel2i interface to expose the CARDs in the absence of PAMP RNA.

**Figure 2.**
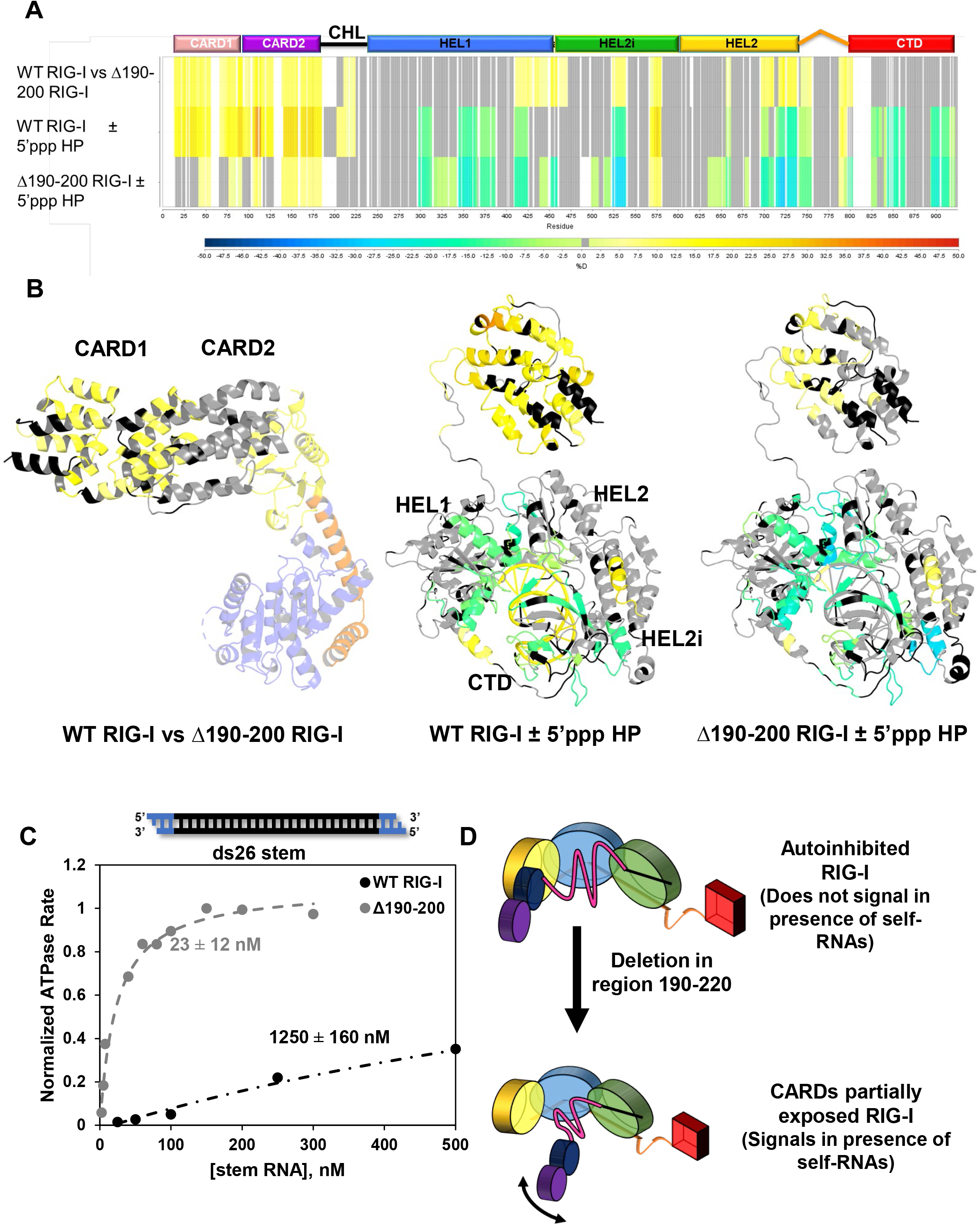
Deletions in the CHL cause partial CARDs exposure and promote high affinity stem RNA binding activity to RIG-I. (A) Hydrogen-Deuterium Exchange Mass Spectroscopy (HDX) heatmap of three comparisons: WT RIG-I vs Δ190-200 RIG-I without RNA; WT RIG-I with and without 5’triphosphate (5’ppp) RNA, a RIG-I PAMP; and Δ190-200 RIG-I with and without 5’ppp RNA. The bar below indicates % deuteration of a given peptide region. White spaces indicate no sequence coverage, and grey represents regions in which the coverage was nonsignificant. (B) Data from (A) modeled onto autoinhibited RIG-I (PDB ID: 4A2W) or activated RIG-I, which is a composite model of two crystal structures (PDB IDs: 5E3H and 4NQK). In the autoinhibited structure, only CARDs and Hel2i are colored according to the HDX results. (C) RIG-I binding to ds26 stem RNA. The ATPase activity of WT RIG-I and Δ190-200 RIG-I was measured at increasing concentration of ds26 stem RNA. In the stem RNA cartoon, black bars indicate RNA region, blue bars indicate DNA region. The DNA overhangs block RIG-I CTD binding at each end. The binding curves were fit using a hyperbola (Equation 5), and the K_D, app_ for each RIG-I construct is reported. (D) Model showing the consequences of CHL deletion. Small deletions in the CHL disrupt both the CHL positioning and the autoinhibitory CARD2:Hel2i interface, causing aberrant CARDs exposure and stem RNA binding. The coloring of the helicase subdomains, CTD, and CARDs is the same as in Figure 1.

We carried out differential HDX-MS analysis of WT and Δ190-200 RIG-I with and without RNA (5’ppp blunt-ended ds10 hairpin) to determine if the CHL deletion had affected RNA binding (**Figure EV2**). We observed a similar degree of protection in the RNA binding regions of the WT and Δ190-200 RIG-I, indicating that CHL deletion does not impair PAMP RNA binding (**Figures 2A and 2B**). Comparison of HDX-MS with and without RNA shows that RNA binding exposes the CARDs in WT RIG-I to a higher degree than in Δ190-200 RIG-I. These results indicate that the CARDs in Δ190-200 RIG-I are partially exposed in the absence of RNA.

To test the second possibility that the CHL deletions impair RNA discrimination, we compared the RNA binding affinities of WT and Δ190-200 RIG-I for a stem RNA mimic. The ds26 stem contains a dsRNA stem region and dsDNA ends, which block RIG-I from end binding, forcing it to only bind internally to the dsRNA region. Thus, the ds26 stem mimics secondary stem structures in self RNAs. The RNA K_D,app_ values were estimated from the binding curves that monitored RIG-I’s RNA-dependent ATPase activity as a function of increasing RNA. WT RIG-I binds the ds26 stem with a weak affinity (K_D,app_ ~1250 nM), as expected (**Figure 2C**). Strikingly, Δ190-200 RIG-I binds the ds26 stem with a 50-fold higher affinity (K_D,app_ of ~20 nM). These results show that disrupting the CHL region dysregulates RNA binding, increasing RIG-I’s affinity for non-specific RNAs.

Thus, the hyperactive signaling responses of the CHL deletion mutants can be explained by its destabilized CARD2:Hel2i interface and dysregulated RNA discrimination mechanism (**Figure 2D**). Thus, CHL is a crucial regulatory region that RIG-I uses to control its aberrant signaling activity in the absence of PAMP RNA.

### The negative charges in the CHL are essential in preventing PAMP-independent RIG-I signaling

Sequence analysis of RIG-I homologs from bony fish to mammals shows that the CHL sequence is poorly conserved compared to the high conservation seen in the helicase domain sequence (**Figure 3A**). Interestingly, CHL is strikingly rich in negatively charged amino acids. Between residues, 186-220 of the human CHL, 11 of the 35 amino acids are either aspartate or glutamate, with only three lysine residues across the CHL. The entire CHL region from 186-241 has a theoretical pI of 3.73. Despite poor sequence conservation, these negative charges in the CHL are well conserved in the RIG-I homologs. This pattern extends to RIG-I-like receptor MDA5, whose larger CHL region has a similar conserved, negative charge pattern despite poor overall sequence conservation (**Figure EV3**). In MDA5, however, electronegative residues are more evenly distributed across the CHL.

**Figure 3.**
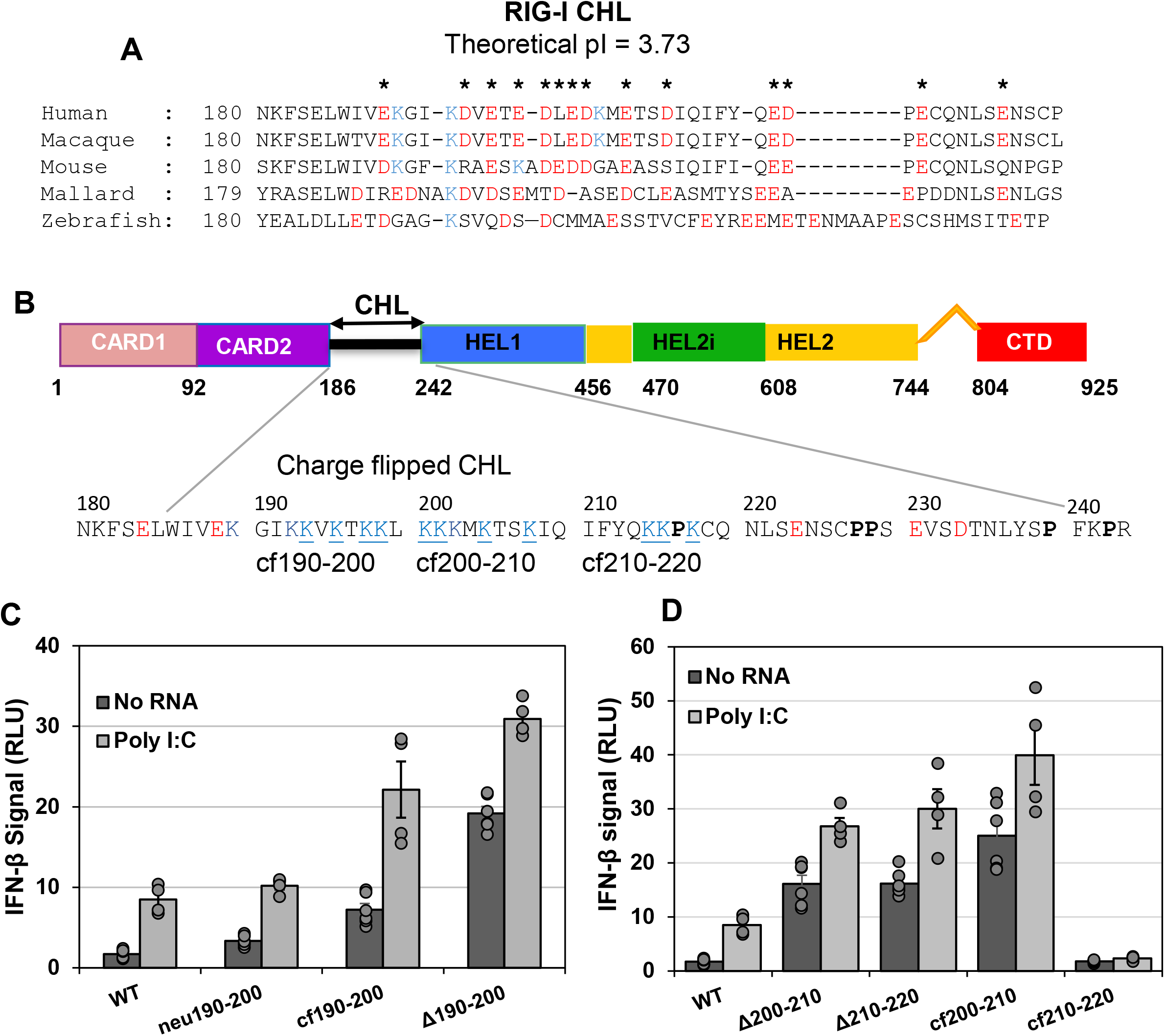
The CHL’s regulatory ability is predicated on its exceptional negatively charged amino acids. (A) Alignment of RIG-I homolog sequences spanning human to bony fish from residues 180-228 shows the conserved negatively charged amino acids in the CHL of all species. Note, this CHL region 190-220 was identified in Figure 1 as important for regulation. All glutamate and aspartate residues are colored red, and lysines colored blue. Asterisks indicate when a negative charge, either glutamate or aspartate, is conserved at that position in 3 out of 5 homologs tested. (B) RIG-I domain schematic with the sequence of CHL highlighted. The sequences of the charge flip mutants (cf190-200, cf200-210, cf210-220) are shown. (C) Cell signaling assays, focused on the 190-200 CHL region, were performed either with or without Poly I:C transfection. Region 190-200 in Neu190-200 was replaced with a glycine-threonine-serine repeat. (D) Same as (C), except focusing on regions 200-210 and 210-220. Note the WT RIG-I results are shared between (C) and (D).

We systematically introduced clusters of neutral or positively charged amino acids in the 190-220 CHL region to determine if CHL’s negatively charged regions are critical in suppressing aberrant RIG-I signaling activity. To make a neutral 190-200 linker (neu190-200), we replaced this segment with a glycine-serine-threonine repeat. To create a positively charged linker, we changed all the aspartate and glutamate residues to lysine in the 190-200 region (cf190-200) (**Figure 3B**). Changing the acidic 190-200 linker to a neutrally charged one increased the PAMP-independent signaling activity by 2-fold, charge flipping increased signaling by 4-fold, and deletion by 10-fold (**Figure 3C**). Interestingly, charge flipping the 200-210 segment (cf200-210) increased the signaling response by 15-fold. The cf210-220 mutant was not expressed for unknown reasons and did not show any signaling responses (**Figure 3D**). Addition of Poly I:C showed the typical increase in signaling activity in all mutants, except for cf210-220, indicating that CHL charge inversion does not interfere with RIG-I’s normal response to PAMP RNA.

The above results show that CHL uses the negatively charged residues in the 190-210 region to suppress RIG-I’s aberrant signaling activity in the absence of PAMP RNA.

### CHL’s negative charges block the initial binding of stem RNA to the helicase domain

Studies above show that CHL plays a crucial role in stabilizing the CARD2:Hel2i interface and regulating RNA binding into the helicase domain, raising many questions about its mechanism of action. Does CHL directly regulate RNA binding or its activity depends on maintaining the CARD2:Hel2i autoinhibitory interface? Are CHL’s negative charges involved in RNA regulation? To answer these questions, we created four RIG-I constructs: WT RIG-I, CHL-Hel-CTD (lacks CARDs but retains CHL), Hel-CTD (lacks CARDs and CHL), and CHL-Hel-CTD cf190-210 (negative residues between 190-210 mutated into lysines) (**Figure 4A**), and tested each protein’s ability to bind the ds26 stem RNA (**Figure 4B**). To measure stem RNA binding in the absence of ATP, we used fluorescence-based titration experiments using a Cy3 fluorophore-labeled ds26 stem. Note that stem RNA has no affinity for the RIG-I’s CTD (Devarkar *et al*., 2018); hence, any RNA binding observed here would be at the helicase domain.

**Figure 4:**
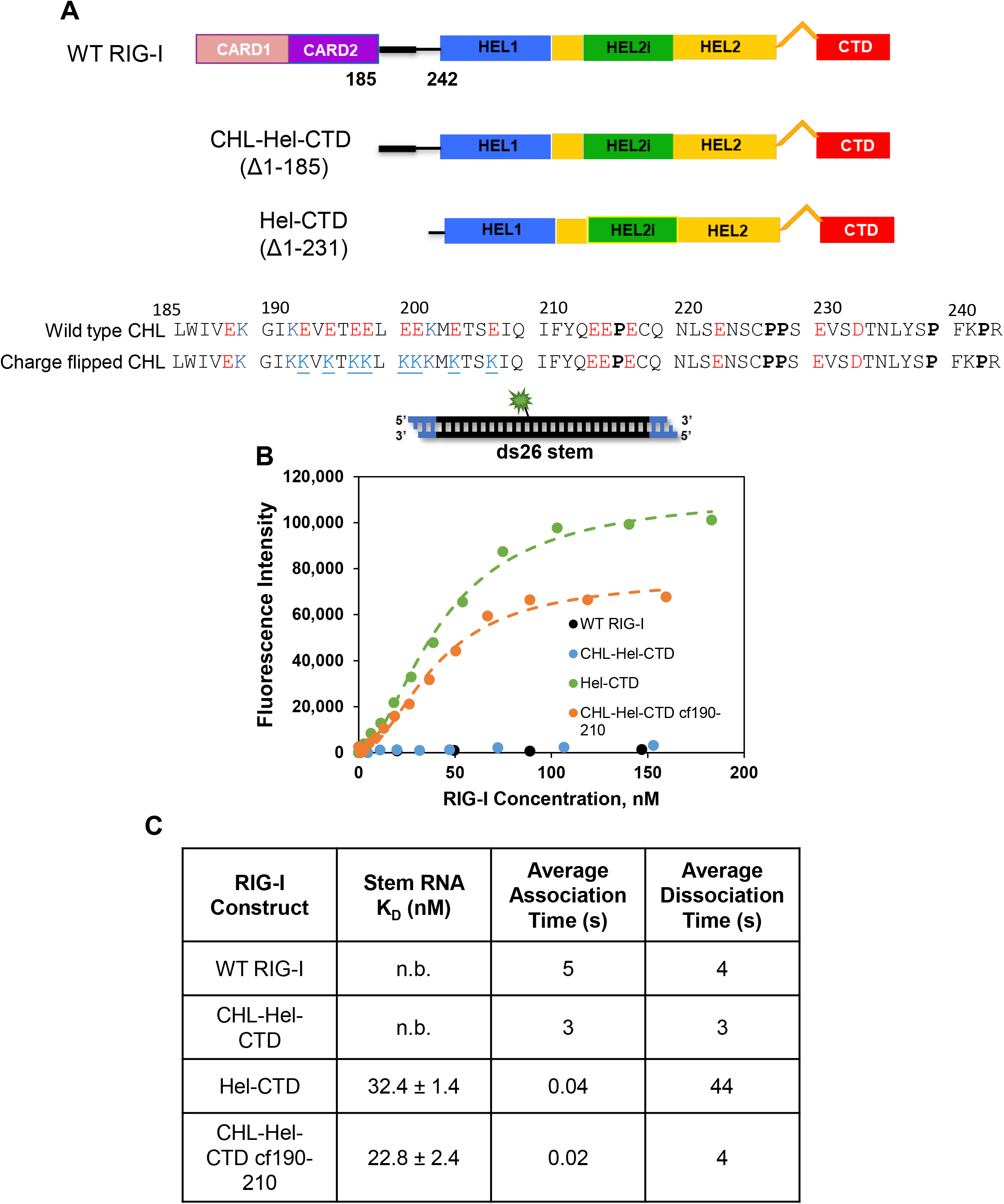
RIG-I’s CHL directly regulates RNA binding through its negative charge. **(A)** Schematics of the tested RIG-I constructs. The 190-210 CHL region identified as being important for regulating signaling activity is highlighted by a thicker black bar. The CHL sequence is highlighted with negative amino acids in red and positive ones in blue. The negatively charged amino acids in the 190-210 region were mutated to lysines to create the CHL-Hel-CTD cf190-210 construct. This change resulted in a pI change in the CHL from to 3.7 to 9.5. **(B)** Fluorescence intensity changes upon binding of various RIG-I constructs to a ds26 stem RNA with an internal Cy3 fluorphore at the 13^th^ position. The RNA binding data of Hel-CTD and CHL-Hel-CTD cf190-210 were fit using a Hill equation (Hel-CTD, n = 1.8 ± 0.2; CHL-Hel-CTD cf190-210 n = 1.9 ± 0.1). **(C)** The Table lists the estimated stem RNA K_D_ values of each RIG-I construct. No binding or n. b. indicates that stem RNA binding was not observed within 400 nM of titrated RIG-I. The average stem RNA association and dissociation times of the various RIG-I constructs were estimated from the stopped-flow experiments shown in Figure EV4 and Table S1.

As expected, the ds26 stem showed undetectable binding to WT RIG-I and high affinity binding to Hel-CTD (K_D_ of 32 nM) (**Figures 4C and 4D**). We reasoned that if CHL has no independent role in regulating RNA binding, then CHL-Hel-CTD will bind the ds26 stem similar to the Hel-CTD. Instead, we observed undetectable binding of the ds26 stem to CHL-Hel-CTD, like the WT RIG-I. Thus, CHL plays a direct role in inhibiting stem RNA binding into the helicase domain. Interestingly, the charge flip mutant bound stem RNA with nM K_D_ value (23 nM), like the Hel-CTD. Thus, CHL uses its negatively charged regions to antagonize stem RNA binding into the helicase domain.

To understand the mechanism of RNA inhibition by the CHL, we measured the ds26 stem association and dissociation rates using stopped-flow experiments. The kinetics of RIG-I binding to with Cy3 labeled ds26 stem (**Figure EV4**) were analyzed to obtain the association rates (**Table S1**). To measure the dissociation rate, a preformed complex of Cy3 labeled ds26 stem with RIG-I was chased with an excess amount of unlabeled competitor RNA. The time-dependent decrease in fluorescence intensities provided the dissociation rates (**Table S1**). Our data show that that ds26 stem binds to CHL-Hel-CTD and WT RIG-I in 3-5 seconds, but under similar conditions, the stem RNA binds to Hel-CTD and the charge inverted CHL-Hel-CTD cf190-210 more in 20 to 40 milliseconds, hence 125 to 250-fold faster (**Figure 4C**). These results clearly show that CHL kinetically inhibits stem RNA binding by blocking its initial interactions with the helicase domain. Furthermore, the kinetic data shows that CHL on its own can kinetically block RNA binding, and the mechanism of blocking involves its negatively charged regions.

To determine if CHL plays a role in RNA dissociation, we compared the lifetimes of ds26 stem on the various RIG-I constructs (**Figure 4C**). The ds26 stem has a relatively long, ~ 40s, lifetime on the Hel-CTD, but the lifetimes of RNA bound to constructs containing the CHL region is ~10-times shorter. These data indicate that CHL destabilizes the stem RNA complex. Strangely, the stem RNA lifetime on the charge flipped CHL-Hel-CTD was similar to CHL-Hel-CTD. Thus, the 190-210 CHL region is not involved in the mechanism of destabilizing stem RNA binding. This function might be performed by the 210-240 CHL region, which remained WT-like in the charged flipped mutant.

### The CHL and CARD2:Hel2i interface establish a tunable gating mechanism for RNA discrimination

Is CHL a general RNA competitor or does it selectively compete with non-specific RNAs not recruited by the CTD? To address this question, we created two new non-PAMP ligands (ds27 with 2-nt 5’-overhang or 3’-overhang), that unlike ds26 stem, contain RNA ends with varying affinities for the CTD. Previous studies show that the 5’ovg RNA end binds CTD with a ~4-fold weaker affinity than the 3’ ovg RNA end (Ramanathan *et al*., 2016). The different affinities enabled us to address the question of RNA selectivity. The ATPase assay was used to measure the RNA K_D,app_ values because having ATP mimics physiological conditions and measuring ATPase activity reports on RNA binding to the helicase domain.

As shown previously (Devarkar *et al*., 2018), the presence of ATP stabilizes the binding of stem RNA (**Figure 5E**). With ATP present, we could measure the K_D,app_ of ds26 stem complex with WT RIG-I (1250 nM) and CHL-Hel-CTD (26 nM), which were not measurable in the absence of ATP due to weak affinity (**Figure 4C**). The contribution of CHL alone to RNA inhibition was determined by comparing the RNA K_D,app_ values of CHL-Hel-CTD to Hel-CTD. The results show that CHL weakens the binding of all three non-PAMP RNAs by 11-15 fold, irrespective of their RNA end modification (**Figures 5C-E** and **Table S2)**. The uniform weakening of non-PAMP RNA binding indicates that CHL is a general RNA competitor. When the negatively charged residues in the 190-210 CHL region were mutated to lysines, CHL lost the ability to inhibit RNA binding and behaved like the Hel-CTD, binding all non-PAMP RNAs with a high affinity. Thus, the negative charges in CHL are essential for RNA competition. The RNA must displace the CHL to bind into the helicase domain and the energy penalty to do so is 1.4 to 1.6 kcal/mol (**Table S3**).

**Figure 5.**
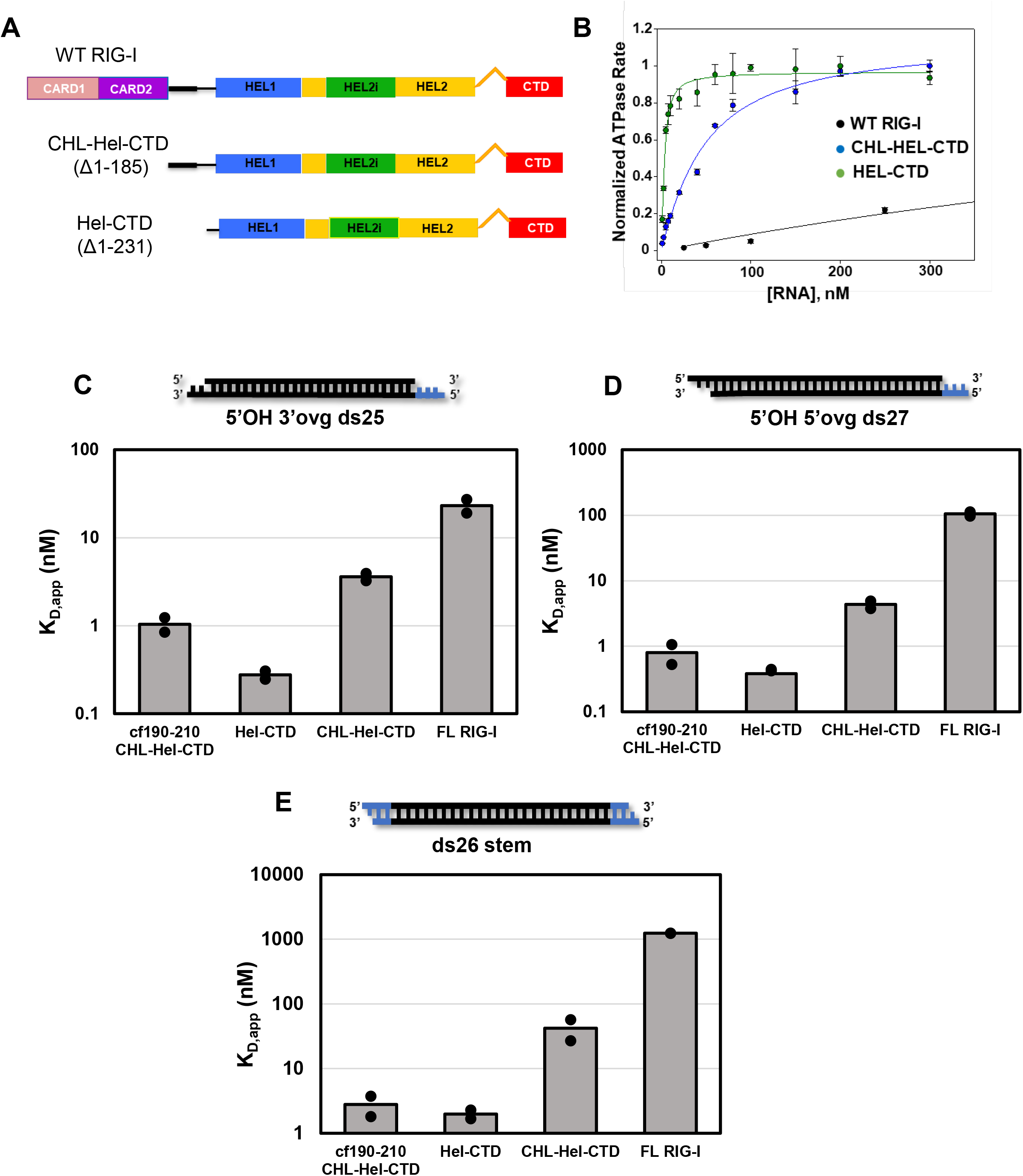
Contribution of CHL and CARD2:Hel2i interface to RNA gating and discrimination. (A) Schematic of purified RIG-I constructs, as in Figure 4. (B) Example of the RNA titration experiments monitoring RIG-I’s ATPase activity as a function of increasing concentration of ds26 stem RNA. Individual points represent relative ATPase rates determined from a time course of ATPase reaction. Standard errors of the fit are shown. The Hel-CTD titrations were fit using a quadratic equation (Equation 6), while all other RNA tested were fit using a hyperbola (Equation 5) to obtain the K_D, app_ values. (C) - (E) Bar charts comparing the K_D, app_ values of (C) 5’OH 3’ovg ds25, (D) 5’OH 5’ovg ds27, and (E) ds26 stem RNA for each of the RIG-I constructs. Note the logarithmic y-axis. Data from two independent experiments are shown.

In the autoinhibited conformation of RIG-I, the CARDs are sequestered by Hel2i through the CARD2:Hel2i interactions, which localizes CHL more closely near the helicase domain (**Figure 1A**). Hence, we expect the autoinhibited CARDs to enhance CHL’s RNA gating function. Indeed, △190-200 RIG-I with a disrupted CARD2:Hel2i interface behaves like CHL-Hel-CTD (**Figure EV5**). In contrast, CHL linked to the CARDs sequestered by the CARD2:Hel2i interface behave differently by more effectively inhibiting non-PAMP RNA binding. Interestingly, RNAs that bind CTD more weakly are blocked more effectively than those binding more tightly. Hence, stem RNA with no affinity for the CTD was inhibited by 700-fold, 5’ ovg RNA by 240-fold, and 3’ ovg RNA by 60-fold. Compared to the energy needed to displace CHL on its own, breaking the CHL and CARD2:Hel2i interactions requires a net free energy of 2.4 to 4 kcal/mol. The range arises from the different energetic contribution of the CTD:RNA end interactions. The model shown in **Figure 6,** and discussed below, explains how CHL and CARD2:Hel2i work at different steps in the RNA binding pathway to selectively block RNAs that are not chosen by the CTD while enabling the CTD-bound RNAs to form a stable complex in the activated RIG-I state.

**Figure 6.**
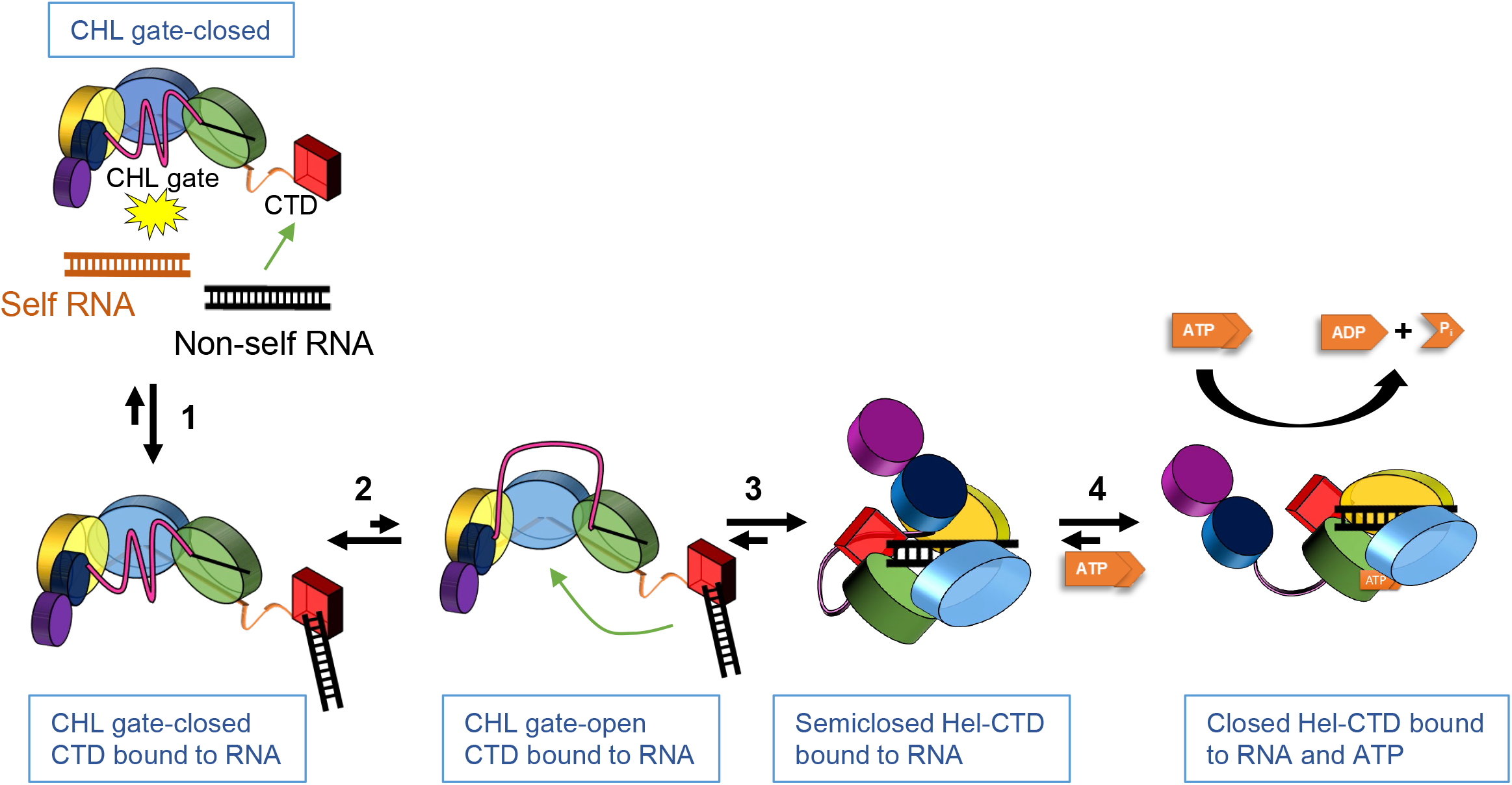
Model showing the roles of the CHL gate and CARD2:Hel2i interface in RNA proofreading. In the autoinhibited RIG-I, the helicase subdomains exist in an open conformation with CARD2 interacting with Hel2i and the CHL gate shielding the helicase. The coloring of the helicase subdomains, CTD, and CARDs is the same as in Figure 1. The CHL gate in the closed-state effectively blocks stem structures in self RNAs from binding directly into the helicase domain. The CTD, however, is free to recognize and bind appropriate RNA ends (step 1). To load the CTD-bound RNA into the helicase domain, the CHL gate must open (step 2). Once the CTD bound RNA makes initial interactions with the helicase domain, the induced conformational change converts the open helicase subdomains to a semiclosed conformation (step 3). In the semiclosed conformation, the CARD2:Hel2i interface is intact, hence the RNA cannot fully engage with the helicase subdomains. We assume that the semiclosed helicase intermediate is not competent in ATP hydrolysis. A favorable binding energy of the RNA with the CTD and helicase is necessary to break the CARD2:Hel2i interface and drive the helicase closing conformational change (step 4). ATP binding facilitates step 4. The helicase in the closed conformation can undergo multiple ATPase turnover as long as RNA remains bound.

## Discussion

This study has identified a previously unknown RNA proofreading mechanism that RIG-I uses to regulate its helicase domain activities. In RIG, the C-terminal domain (CTD) serves as the RNA sensor and the helicase domain as an actuator, switching RIG-I from the autoinhibited state to the activated state in response to appropriate RNAs. The helicase must bind RNAs chosen by the CTD and not to self RNAs present in the cytoplasm to ensure that specific RNAs activate the autoinhibited CARDs. Our study reveals the regulatory mechanism that controls RNA binding into the helicase domain. We show that the CARDs-Helicase Linker (CHL), previously considered a passive linker, is an essential part of this regulatory mechanism that stabilizes the CARD2:Hel2i interface while electrostatically gating the helicase domain from non-specific RNA binding events.

Sequence analysis shows that CHL is intrinsically disordered and highly negatively charged. Such intrinsically disordered linkers are increasingly recognized to have regulatory functions (van der Lee *et al*, 2014; Vuzman & Levy, 2012; Wright & Dyson, 2015). Consistent with its dynamic nature, the 187-244 CHL region was not resolved in the autoinhibited duck RIG-I structure (Kowalinski *et al*., 2011). However, the structure shows the missing 57 amino acids, spanning the 40 Å distance between CARD2 and Hel1, are located close to the helicase domain. Interestingly, the AlphaFold structure prediction also places the CHL into the central helicase binding channel (Jumper *et al*, 2021). Being long and disordered, the CHL can exclude a significant volume near the helicase domain to block RNA binding in the autoinhibited conformation of RIG-I (**Figure 1**). Our RNA binding studies confirmed this hypothesis and showed that CHL kinetically competes with RNAs entering the helicase domain. The ability of the CHL to kinetically compete with incoming RNAs was compromised drastically when the negatively charged amino acids in the 190-210 region were mutated to lysines. Thus, CHL electrostatically gates the helicase to control RNA binding.

In addition to serving as an RNA gate, the negatively charged CHL also stabilizes the CARD2:Hel2i interface. The HDX-MS analysis of the △190-200 RIG-I indicated that this small deletion destabilizes the CARD2:Hel2i interface and spontaneously exposes the CARDs. The 190-200 deletion also dysregulated RNA selection, with the mutant showing higher affinities for non-PAMP RNAs, like the stem RNA and 3’ovg and 5’ovg RNAs, compared to WT RIG-I. The destabilized CARD2:Hel2i interface and high affinities for self RNAs result in hyperactive immune response in the absence of PAMP RNA. Therefore, CHL regulates the helicase domain activities in multiple ways that are essential in keeping RIG-I in a signaling-silent state in the absence of viral infection. The CHL region is not unique to the RIG-I. MDA5, the other member of the RIG-I family, also contains a negatively charged CHL region between its CARDs and the helicase domain, close to twice the size of RIG-I CHL (**Figure EV3**). It will be interesting to determine if MDA5’s CHL plays a similar role in regulating RNA binding and CARDs activation.

The CHL on its own is a general RNA competitor, but when CHL is linked to the CARDs stabilized by the CARD2:Hel2i interface, the two work together to establish a gating mechanism that preferentially allows CTD-bound RNAs to bind into the helicase domain while excluding non-specific RNAs. The RNA binding mechanism proposed in **Figure 6** explains how the CHL gate and CARD2:Hel2i interface discriminate between self and non-self at different steps in the RNA binding pathway. In the autoinhibited RIG-I, the CHL gate blocks RNAs from directly binding into the helicase domain. As our previous studies showed (Devarkar *et al*., 2018), the CTD is free to survey RNA ends and bind to PAMP RNA (step 1). However, the CTD-bound RNAs cannot load into the helicase domain until the CHL gate assumes an open-gate conformation. The CHL gate being dynamic likely exists in the gate-closed and gate-open states (step 2). Those RNAs that do not bind to the CTD, like most self RNAs, have a low probability of accessing the gate-open conformation from the solution. The likelihood of gate opening coinciding with the RNA collision is low. On the other hand, CTD-bound RNAs are near the helicase domain and hence can access the gate-open state with a higher probability. Especially, RNAs with a longer lifetime on the CTD, like the PAMP RNAs, have a greater chance of binding into the helicase domain than those with lower affinities like the overhang RNAs. By preferentially loading CTD-bound RNA, the CHL gating mechanism effectively antagonizes self RNAs. In this manner, the CHL gate discriminates between self and non-self at the very first step of RNA binding.

Once the RNA interacts with the helicase, the induced conformational change converts the helicase from an open state to a semiclosed conformation (step 3). Structural studies of RIG-I provides evidence for open, semiclosed, and closed forms of the helicase domain in RIG-I (Civril *et al*., 2011; Devarkar *et al*., 2016; Jiang *et al*., 2011; Kowalinski *et al*., 2011; Luo *et al*., 2011; Luo *et al*., 2012). In the semiclosed state, the CARD2:Hel2i interface is intact and the RNA is not fully engaged with the helicase subdomains. To fully engage the RNA, the helicase must assume a closed conformation by breaking the CARD2:Hel2i interface, facilitated by the free energies of RNA and ATP binding (step 4). This is the stage where the CARD2:Hel2i interface contributes to RNA selectivity. The binding energy of the RNA with the helicase and the CTD provides the energy to break the CARD2:Hel2i interface. Non-specific RNAs that do not interact with the CTD do not have the necessary free energy to break the CARD2:Hel2i interface and activate the CARDs. On the other hand, PAMP RNA ends are strong, providing the necessary amount of free energy to efficiently activate the CARDs. To test the above four-step mechanism with our experimental data, we modeled the RNA binding data of WT RIG-I using the KinTek Explorer software (**Figure EV6**). The above model with an unfavorable gate opening step of 0.01 predicted the expected trend, an unfavorable equilibrium constant of ~0.7 for the helicase closing step with the ds26 stem RNA, and increasingly favorable equilibrium constants of ~3 for the 5’ovg RNA and ~5 for the 3’ovg RNA.

The discovery of the regulatory functions of the intrinsically disordered CHL region represents an exciting, new, and unique opportunity to leverage this region in future RIG-I-based therapeutics. The CHL being more specific than the ubiquitous ATPase pocket, can serve as a new drug target site to modulate RIG-I activity. Compounds that destabilize the CHL, akin to deletions and substitutions, may represent a new class of RIG-I activating molecules that can stabilize the CARDs-released state without directly targeting the CARD2:Hel2i interface.

## ACKNOLWEDGEMENTS

We thank the members of the Patel Lab for their feedback and support throughout these studies. We thank Neel Drain, Arjun Gupta, and Jasper Albers for performing this project’s pilot experiments in their summer high school project. This work was supported by the NIGMS MIRA grant GM118086 (to S.S.P.).

## AUTHOR CONTRIBUTIONS

Conceptualization, B.D.S., S.S. P., S.C.D.; Investigation, B.D.S., S.C.D., J.Z.; Writing – Original Draft, B.D.S., S.S.P.; Writing – Review/Editing: All Authors; Funding Acquisition, S.S.P., P.R.G.

## Materials and Methods

HEK293T cells (Sex: Female) were a gift from Prof. Charles Rice, Rockefeller University USA. This cell line was described and used previously (Devarkar *et al*., 2018; Ramanathan *et al*., 2016), as it is devoid of RIG-I expression and does not respond to transfected double-stranded RNA or poly I:C. Cells were grown at 5% CO_2_ and 37°C, in DMEM with 10% FBS. RIG-I was constitutively expressed in cells by plasmid transfections and were then tested for IFN-β cell signaling response to various transfected RNA ligands.

### IFN-β Reporter cell signaling assays

HEK293T cells were grown in 6-well plates to 60% confluence and cotransfected with firefly luciferase reporter plasmid (pLuc125 / 2.5 μg), Renilla luciferase reporter plasmid (pRL-TK / 500 ng), and a plasmid carrying either the wt RIG-I gene or mutant RIG-I construct under the constitutively active CMV promoter (pcDNA 3.1 / 2 μg). The firefly luciferase gene is under the control of the interferon β promoter, and the Renilla luciferase plasmid is under the control of constitutively active TK promoter. The plasmid transfections were carried out with X-tremeGENE HP DNA Transfection Reagent (Roche). Cells were replated in 96-well plates the next day at 2 × 10^4^ cells/well density and transfected with poly I:C (700 ng in 110 μl volume) using Lyovec transfection reagent (InvivoGen). After 20 hours, the activities of firefly and Renilla luciferases were measured sequentially with the Dual-Luciferase reporter assay (Promega). Data was collected in quadruplicate sets for poly I:C transfections and sextuplicate sets for no RNA transfection, and relative luciferase activities were calculated. Error bars represent the standard error of the mean (SEM). Protein expression was tested using western Blots with a primary α-Myc antibody and β-actin as a normalization control.

### Protein expression and purification

RIG-I was expressed using pET28 SUMO vector in *Escherichia coli* strain Rosetta (DE3) (Novagen). Soluble cell lysate was purified using a Ni^2+^-nitrilotriacetate (QIAGEN) column, followed by Ulp1 protease digestion to remove 6xHis-SUMO tag. It was further purified by hydroxyapatite (CHT-II, Bio-Rad) and heparin sepharose column chromatography (GE Healthcare). Purified protein was dialyzed into 50mM HEPES pH 7.5, 50mM NaCl, 5mM MgCl_2_, 5mM DTT, and 10% glycerol overnight at 4°C, frozen in liquid nitrogen and stored at −80°C (Jiang *et al*., 2011).

### RNA

RNAs were chemically synthesized and HPLC purified by Biosynthesis, Dharmacon, and Trilink BioTechnologies. Synthetic RNAs were purity-analyzed using mass spectrometry and HPLC. Lyophilized RNA was resuspended in 20 mM potassium phosphate buffer 7.0. Concentrations were determined from the A_260_, measured using a NanoDrop spectrophotometer, in 7 M guanidinium HCL, as well as the extinction coefficient for each RNA. RNAs were annealed by mixing complementary ssRNAs in a 1:1.1 ratio (fluorescent:nonfluorescent ratio, where applicable), then heating to 95°C, then finally cooling slowly to 4°C.

### Hydrogen-Deuterium exchange (HDX) detected by mass spectrometry (MS)

Peptides were identified using tandem MS (MS/MS) with an Orbitrap mass spectrometer (Q Exactive, ThermoFisher). Product ion spectra were acquired in data-dependent mode with the top five most abundant ions selected for the product ion analysis per scan event. The MS/MS data files were submitted to Mascot (Matrix Science) for peptide identification. Peptides included in the HDX analysis peptide set had a MASCOT score greater than 20, and the MS/MS spectra were verified by manual inspection. The MASCOT search was repeated against a decoy (reverse) sequence and ambiguous identifications were ruled out and not included in the HDX peptide set.

For HDX-MS analysis, 10 μM of WT/Δ190-200 RIG-I receptors (50 mM HEPES, pH 7.4, 150 mM NaCl, 5% glycerol, 5 mM MgCl_2_, 2 mM DTT) were incubated with the RNA ligand at a 1:1.2 molar ratio for 1 hr (protein:ligand) before the HDX reactions at 4 °C. Five-microliter of protein/protein complex with ligand/peptide was diluted into 20 μl D_2_O (deuterium) in exchange buffer (50 mM HEPES, pH 7.4, 150 mM NaCl, 5 mM MgCl_2_, 2 mM DTT) and incubated for various HDX time points (e.g., 0, 30, 60, 300, 600, 900, 1800 and 3600 s) at 4 °C and quenched with 25 μl of ice-cold 4 M guanidine hydrochloride, 1% trifluoroacetic acid. Dmax samples were incubated in D_2_O in exchange buffer containing 3M guanidine hydrochloride (50 mM HEPES, pH 7.4, 150 mM NaCl, 5 mM MgCl_2_, two mM DTT and 3M guanidine hydrochloride) overnight at room temperature. The samples were immediately placed on dry ice after the quenching reactions until they were injected into the HDX platform. Upon injection, samples were passed through an immobilized pepsin column (2mm × 2cm) at 200 μl min^−1^, and the digested peptides were captured on a 2mm × 1cm C_8_ trap column (Agilent) and desalted. Peptides were separated across a 2.1mm × 5cm C_18_ column (1.9 μm Hypersil Gold, ThermoFisher) with a linear gradient of 4% - 40% CH_3_CN and 0.3% formic acid, over 5 min. Sample handling, protein digestion, and peptide separation were conducted at 4°C. Mass spectrometric data were acquired using an Orbitrap mass spectrometer (Q Exactive, ThermoFisher) with a measured resolving power of 65,000 at *m/z* 400. HDX analyses were performed duplicate or triplicate, with single preparations of each protein ligand complex. The intensity weighted mean m/z centroid value of each peptide envelope was calculated and subsequently converted into a percentage of deuterium incorporation. In the absence of a fully deuterated control, corrections for back-exchange were made by an estimated 70% deuterium recovery, and accounting for the known 80% deuterium content of the deuterium exchange buffer. When comparing the two samples, the perturbation %D is determined by calculating the difference between the two samples. HDX Workbench colors each peptide according to the smooth color gradient HDX perturbation key (D%) shown in each indicated figure. Differences in %D between −5% to 5% are considered non-significant and are colored gray according to the HDX perturbation key. In addition, unpaired t-tests were calculated to detect statistically significant (p<0.05) differences between samples at each time point. At least one time point with a p-value less than 0.05 was present for each peptide in the data set, confirming that the difference was significant.

### Fluorimetric titrations to measure RNA Binding

Fluorescence intensity measurements were carried out at 25°C using FluoroMax-4 spectrofluorimeter (Horiba Jobin Yvon) in Buffer A (50 mM MOPS pH 7.4, 5 mM DTT, 5 mM MgCl_2_, 0.01% Tween20). Cy3-labeled ds26 stem RNA (15 nM) was titrated with increasing concentration of RIG-I proteins, and the change in fluorescence intensity was measured at 570 nm after excitation at 547 nm. The observed fluorescence intensity change (F) from the original fluorescence intensity (F_0_) is proportional to the amount of protein-RNA complex (PR) and modified by a coefficient of complex formation (f_c_). This value was plotted as a function of protein concentration (P) and fitted to Equation 1 to obtain the equilibrium dissociation constant (*K*_D_) and Hill coefficient (*n*). The reported RNA *K*_D_ values were consistently observed in titrations repeated two times.

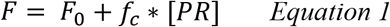

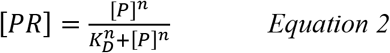

### Stopped-flow experiments to measure RNA association and dissociation rate constants

The RNA dissociation rates were measured at 25°C using a stopped-flow instrument (Auto-SF 120, Kintek Corp, Austin, Tx). A mixture of Cy3-labeled ds26 stem RNA mimic (100 nM) and protein (400 nM) in Buffer A (pre-incubated at 25°C for 10 min) from syringe A was mixed with a 10-fold excess of an RNA trap (unlabeled 5’ppp ds12 hairpin RNA) from syringe B. The Cy3 fluorescence emission was measured using a 570 nm band-pass filter after excitation at 547 nm. The change in fluorescence intensity was plotted as a function of time, and the data were fit to a single exponential or sum of two to three exponentials to estimate the dissociation rates.

The RNA association rates were measured at 25°C using the stopped-flow instrument. A fixed concentration (100 nM) of Cy3-labeled ds26 stem RNA from syringe A was rapidly mixed with the RIG-I construct (250 nM) from syringe B. The fluorescence intensity was plotted as a function of time and fitted a single exponential equation or sum of two to three exponentials to obtain the observed rates of binding.

Fitting association and dissociation kinetics yielded the rate constant (k) and the population fraction (A) associated with each phase. Average association and dissociation times (t) were calculated by taking the inverse of the rate constant and weighting it by population fraction, as shown below in *Equation 3*. Note that the sum of all population fractions for any given fitting will equal 1 (*Equation 4*).

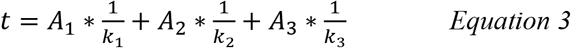

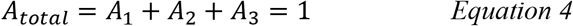

### RNA binding using ATP hydrolysis

ATP hydrolysis activity and calculated RNA binding activity were measured at constant RIG-I (15 nM) and increasing RNA concentration (1 nM – 5 μM) in the presence of 1 mM ATP (spiked with [γ-^32^P] ATP). A time course (0, 20, 40, 60 min) of the ATPase reactions was performed in Buffer A at 25°C. Reactions were stopped at each time point using 4 N formic acid and analyzed by PEI-Cellulose-F TLC (Merck) developed in 0.4 M potassium phosphate buffer (pH 3.4). TLC plates were exposed to a phosphorimager plate, imaged on a Typhoon phosphor-imager, and quantified using the ImageQuant software. The ATPase rate was determined from the slopes of molar [Pi] produced versus time (s). These rates were plotted as a function of RNA concentration and fitted to the following: ATPase rate = *k*_atpase_ x [PR]/[Pt]; where [Pt] is total protein concentration; [PR] is the amount of RIG-I/RNA complex formed and [R] is the RNA concentration being titrated. The hyperbolic equation (Equation 5) or quadratic equation (Equation 6) was used to determine [PR] and estimate the equilibrium dissociation constant (*K_D,app_*) and maximal ATPase rate (*k*_atpase_) of the RIG-I constructs. The energetic contributions of both the CHL and the combination of CHL-CARDs are described in Equations 7, 8, and 9.

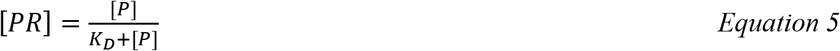

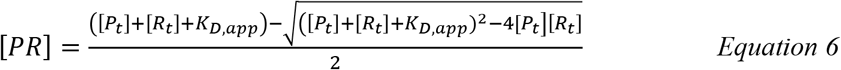

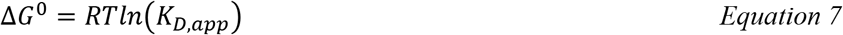

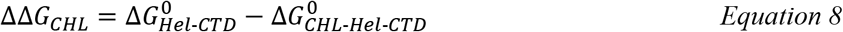

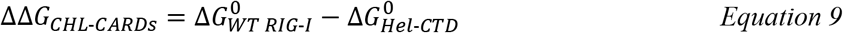

### Quantification and statistical analysis

Luciferase signals in cell-based assays were measured using a luminometer (Tecan) and quantified in Microsoft Excel. The fluorescence intensity titrations, ATP hydrolysis assays, and Hel-CTD titration cell-signaling assays were quantified and fitted using SigmaPlot v11.0 (Systat Software). Stopped-flow kinetics were fitted with Kintek Stop-flow software. All modeling was done with Kintek Explorer 6 software. Associated errors of measurements and fitting and the number of sample sets are noted in the corresponding figure legends.

### KinTek Explorer Data Modeling

The RNA binding data of the WT RIG-I were fit to the model shown in **Figure EV6** using the KinTek Explorer software. In these experiments, we measured the kinetics of Pi formation at increasing RNA concentrations (stem RNA, 3’ovg RNA, and 5’ovg RNA). The rate of Pi formation increases hyperbolically with increasing RNA to provide an apparent K_D,app_ of RNA binding to the helicase domain, reported in **Figure 5**. We normalized the ATPase rates of the three RNAs, and fit the RNA dependency at a fixed concentrations of ATP and RIG-I to experimental conditions of 1 mM and 15 nM, respectivelyThe ADP off rate was assumed to be fast (100 s^−1^). We used the previously reported ATP K_D_ of 5 μM (Devarkar *et al*., 2018), RNA K_D_ for the CTD (Ramanathan *et al*., 2016), and helicase domain RNA K_D_ of ~ 2 μM (Vela *et al*., 2012). Best fits estimated the equilibrium constants of the CHL-gate opening and helicase closing steps.

#### Quantification and Statistical Analysis

Luciferase signals in cell-based signaling assays were measured using a luminometer (Tecan Spark) and quantified in Microsoft Excel. Steady state fluorescence intensity binding assays and ATP hydrolysis-based binding assays were quantified and fitted using SigmaPlot v11.0 (Systat Software). Stopped-flow kinetic traces were fit using the Kintek Stop-flow software. All modeling was done using Kintek Explorer 6 software. Associated errors of the measurements and number of sample sets (n) are noted in the corresponding figure legends.

### Reagents and Tool Table

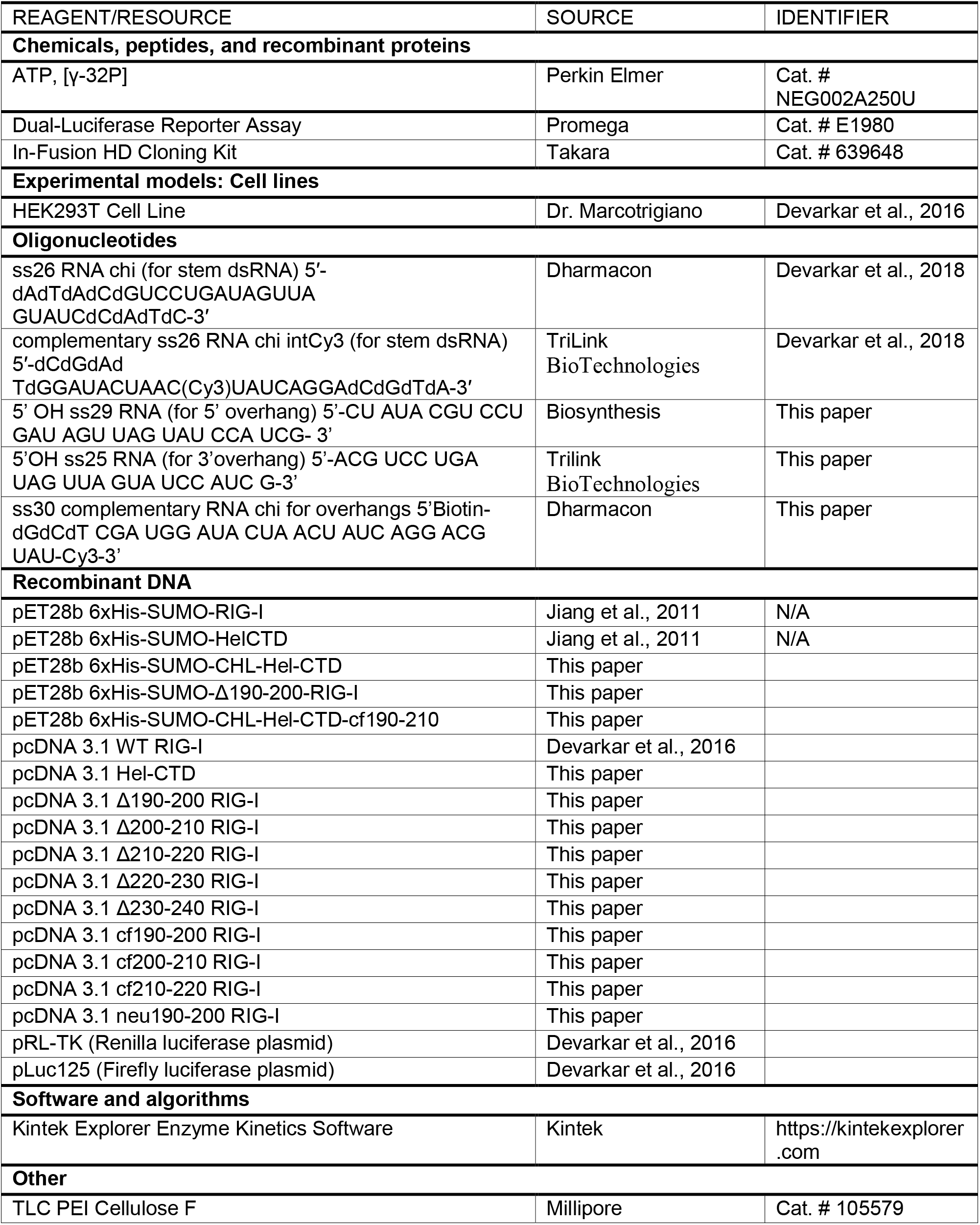

---

**Figure EV1.**
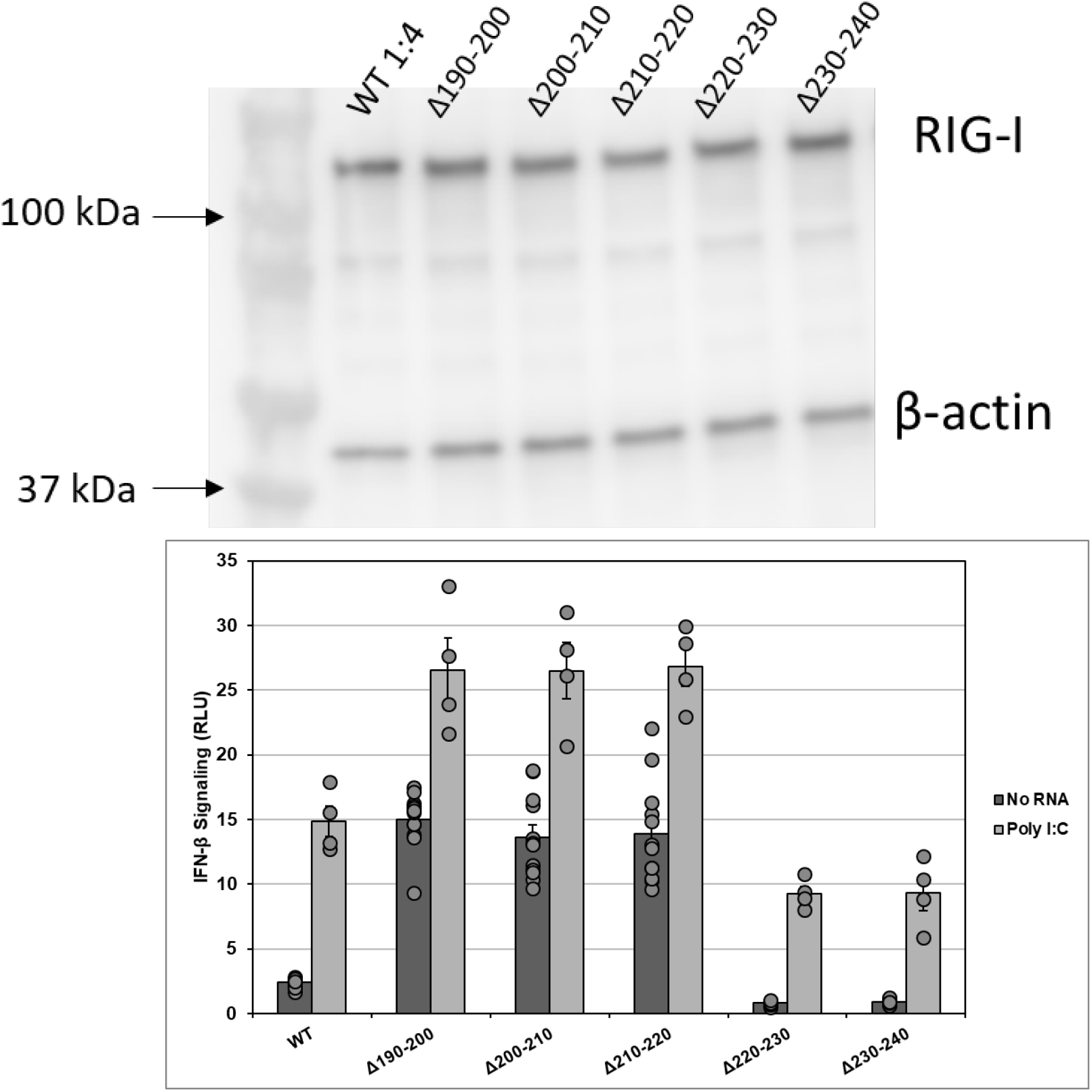
Western Blot of signaling data shown in Figures 1, with respective cell signaling data repeated above each band, to confirm protein expression. In each experiment, pcDNA3.1 myc-tagged RIG-I constructs (approximately 108 kDa) were recognized with a primary α-Mycantibody. β-actin (approximately 42 kDa) was used as a normalization control. (related to Figure 1)

**Figure EV2.**
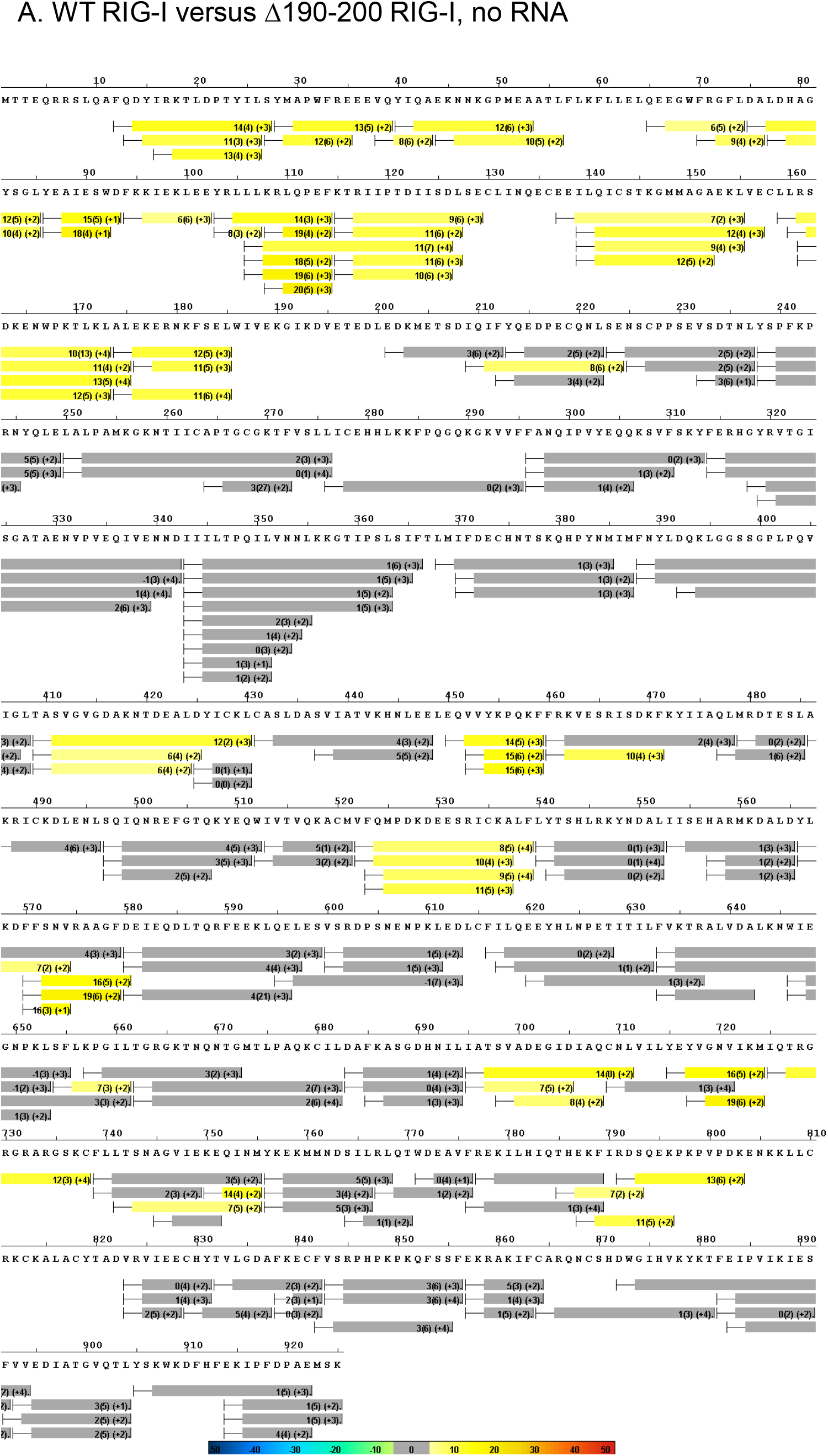
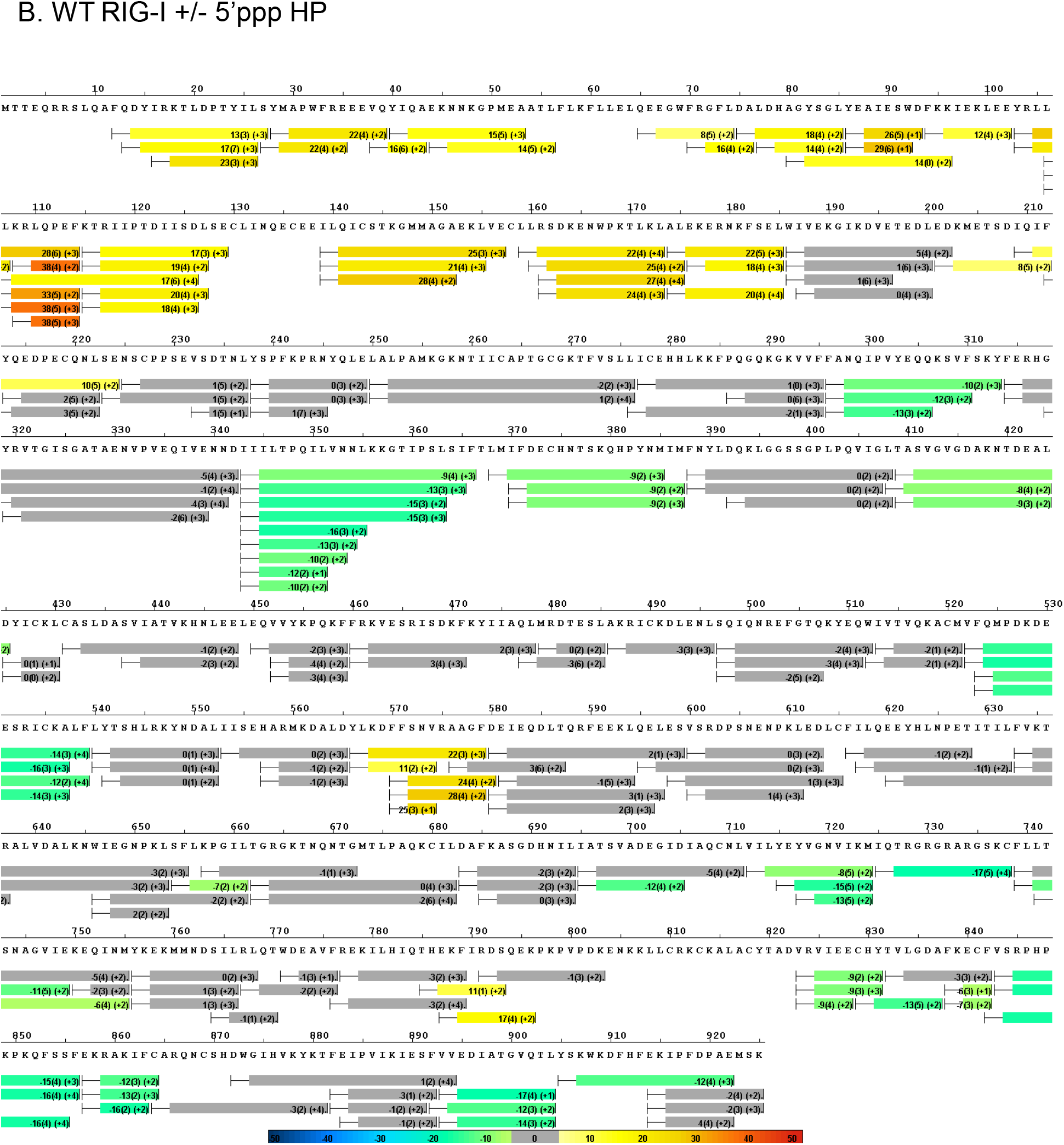
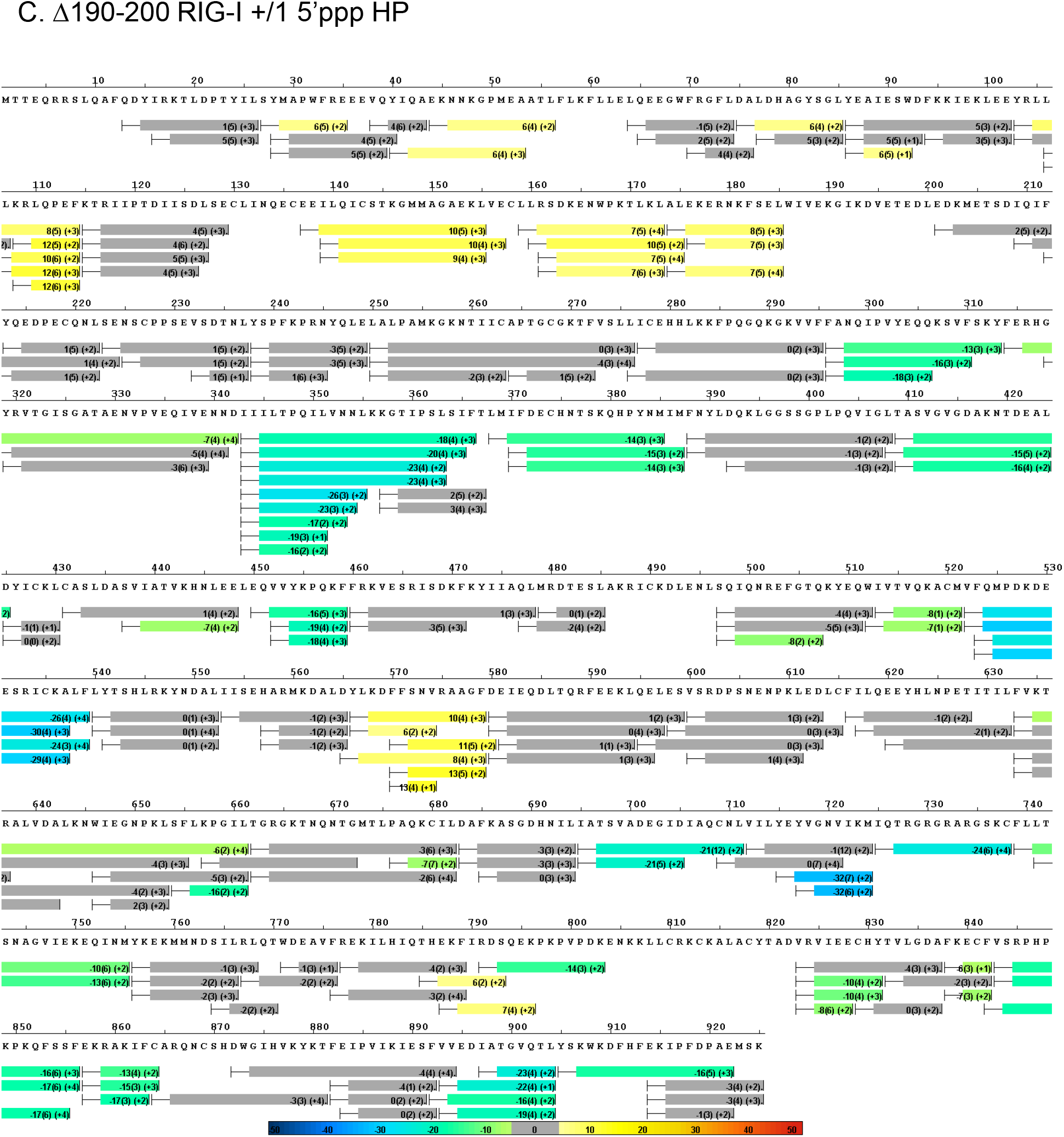
Panels A-C shows the average ΔD2O% ± standard deviation between the two samples across all HDX time points. HDX Workbench colors each peptide according to the smooth color gradient HDX perturbation key shown in each indicated figure. Average ΔD2O% between −5% to 5% are considered non-significant and are colored gray (related to Figure 2).

**Figure EV3.**
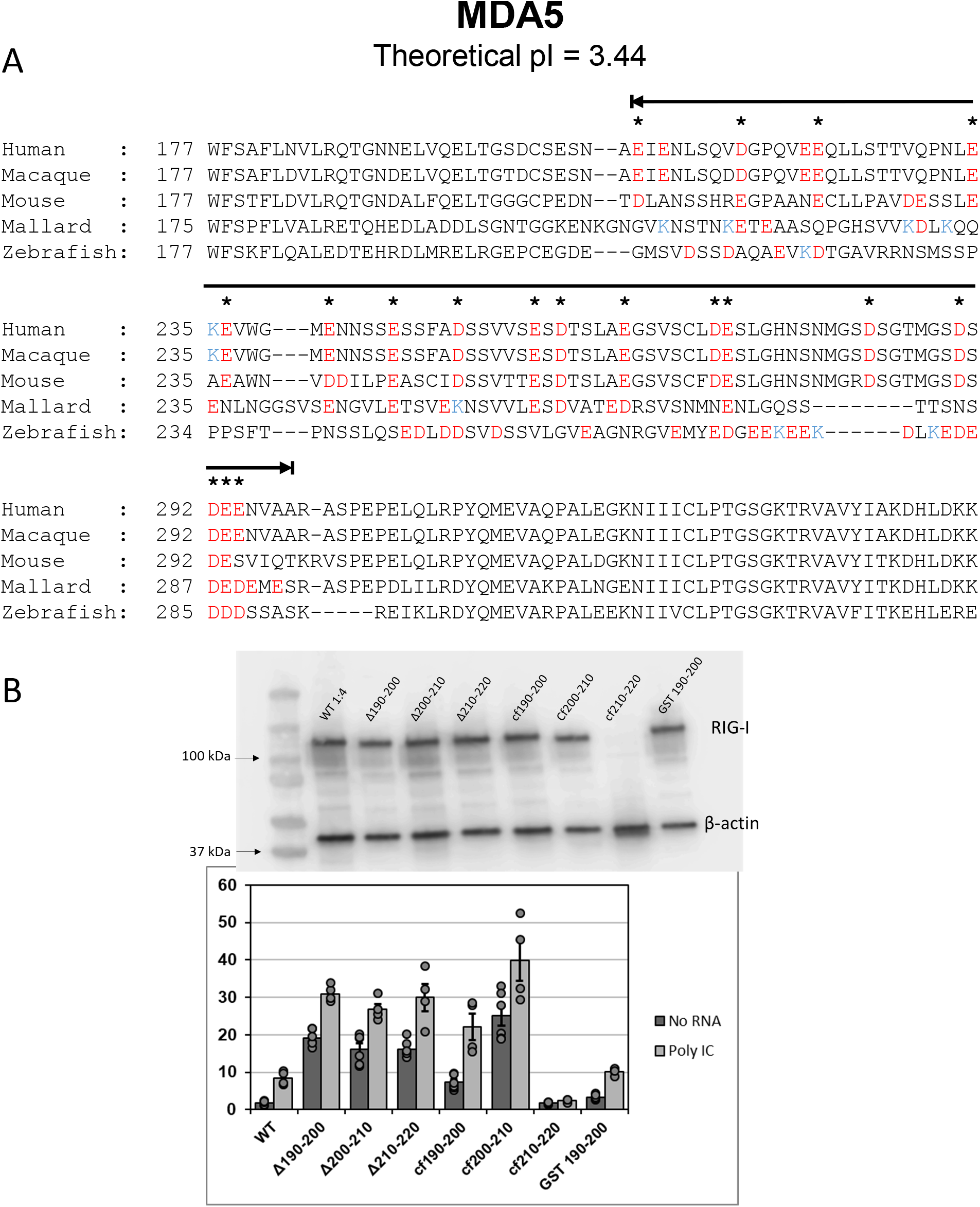
(A) RIG-I Like Receptor MDA5 has an electronegative CARD2-helicase linker region similar to RIG-I. MDA5 sequences from five homologs spanning human to bony fish was aligned and analyzed as in Figure 3A. Black line indicates the MDA5 CHL. (B) Western Blot of signaling data shown in Figures 3, with respective cell signaling data repeated above each band, to confirm protein expression. In each experiment, pcDNA3.1 myc-tagged RIG-I constructs (approximately 108 kDa) were recognized with a primary α-Myc antibody. β-actin (approximately 42 kDa) was used as a normalization control. (related to Figure 3)

**Figure EV4.**
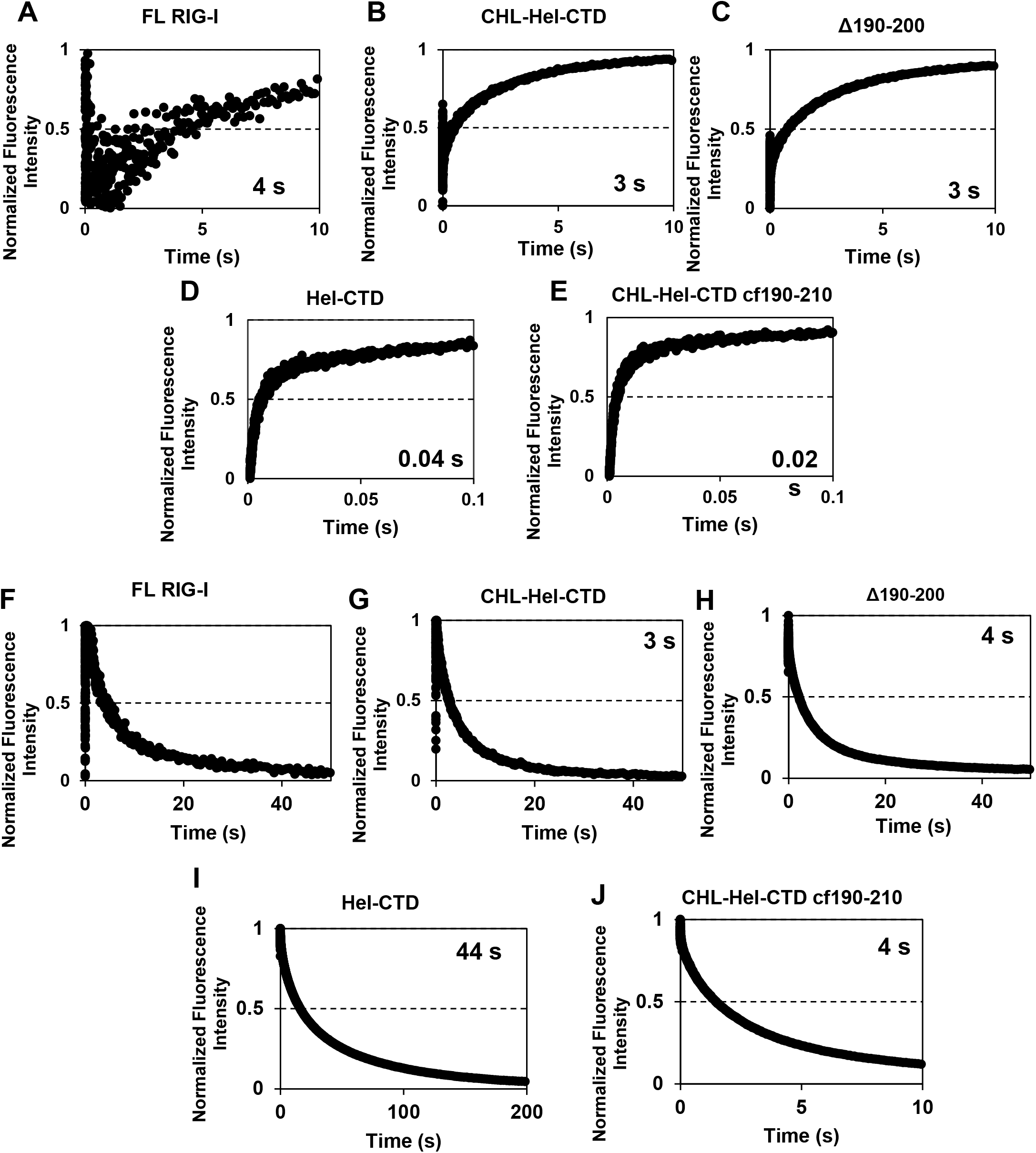
Thumbprint graphs of RIG-I:ds26 stem RNA association (A-E) and dissociation (F-J) kinetics of various RIG-I constructs measured using the stopped-flow experiments described in the Methods. Numbers in each graph are the average association or dissociation times calculated from the amplitude and rate constants shown in Table S2 using Equation 3 (related to Figure 4).

**Figure EV5.**
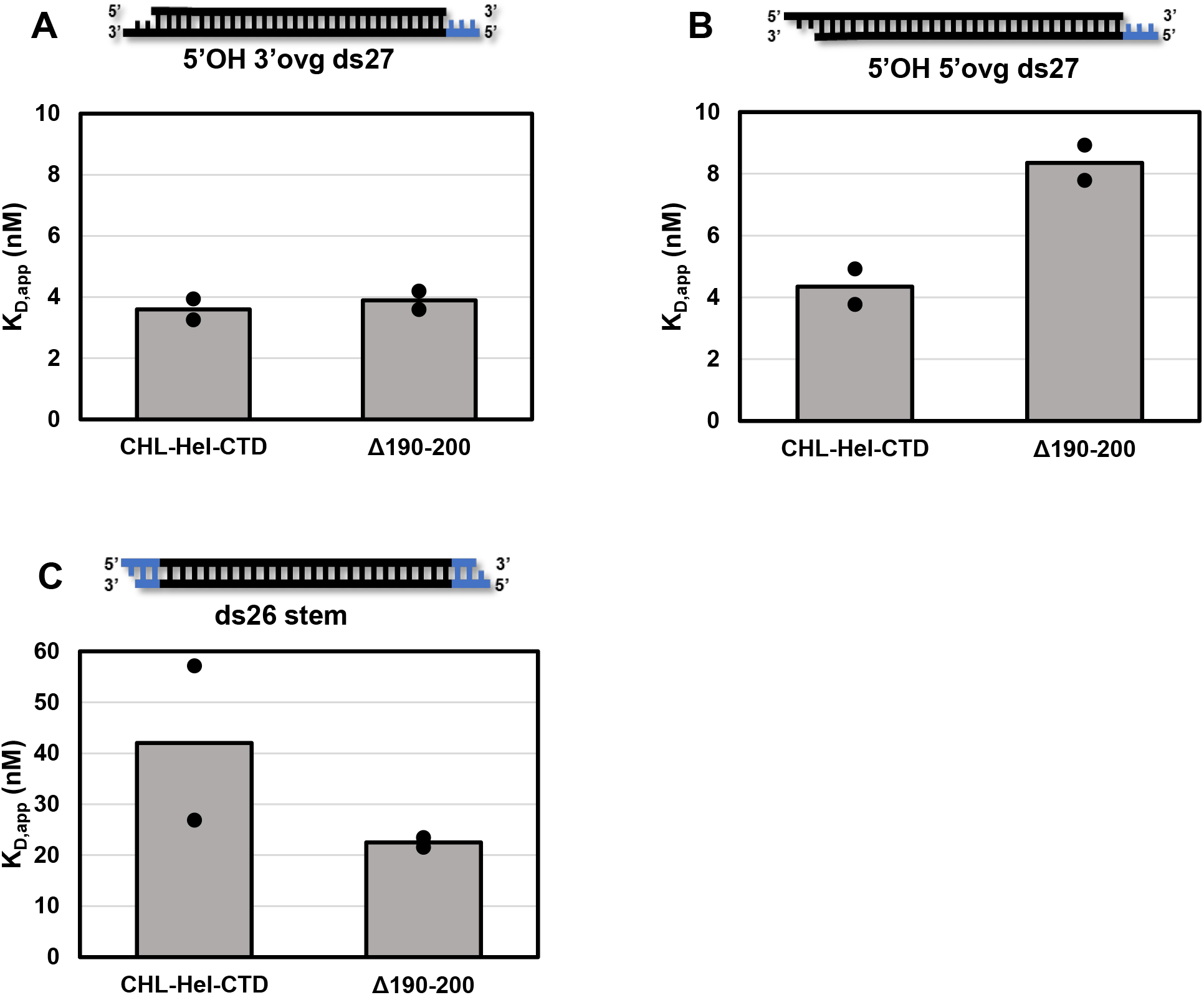
Δ190-200 RIG-I binds RNAs similarly to CHL-Hel-CTD. (A) – (C) Bar charts showing the K_D, app_ from titrations measuring ATPase as a function of RNA concentration for (A) 5’OH 3’ovg ds27, (B) 5’OH 5’ovg ds27, and (C) ds26 stem RNA. Black lines in the cartoon shows the RNA region and blue lines the DNA overhangs added to stem RNA to block CTD binding to the ends. Dots indicate results from two independent RNA binding experiments (related to Figure 5).

**Figure EV6.**
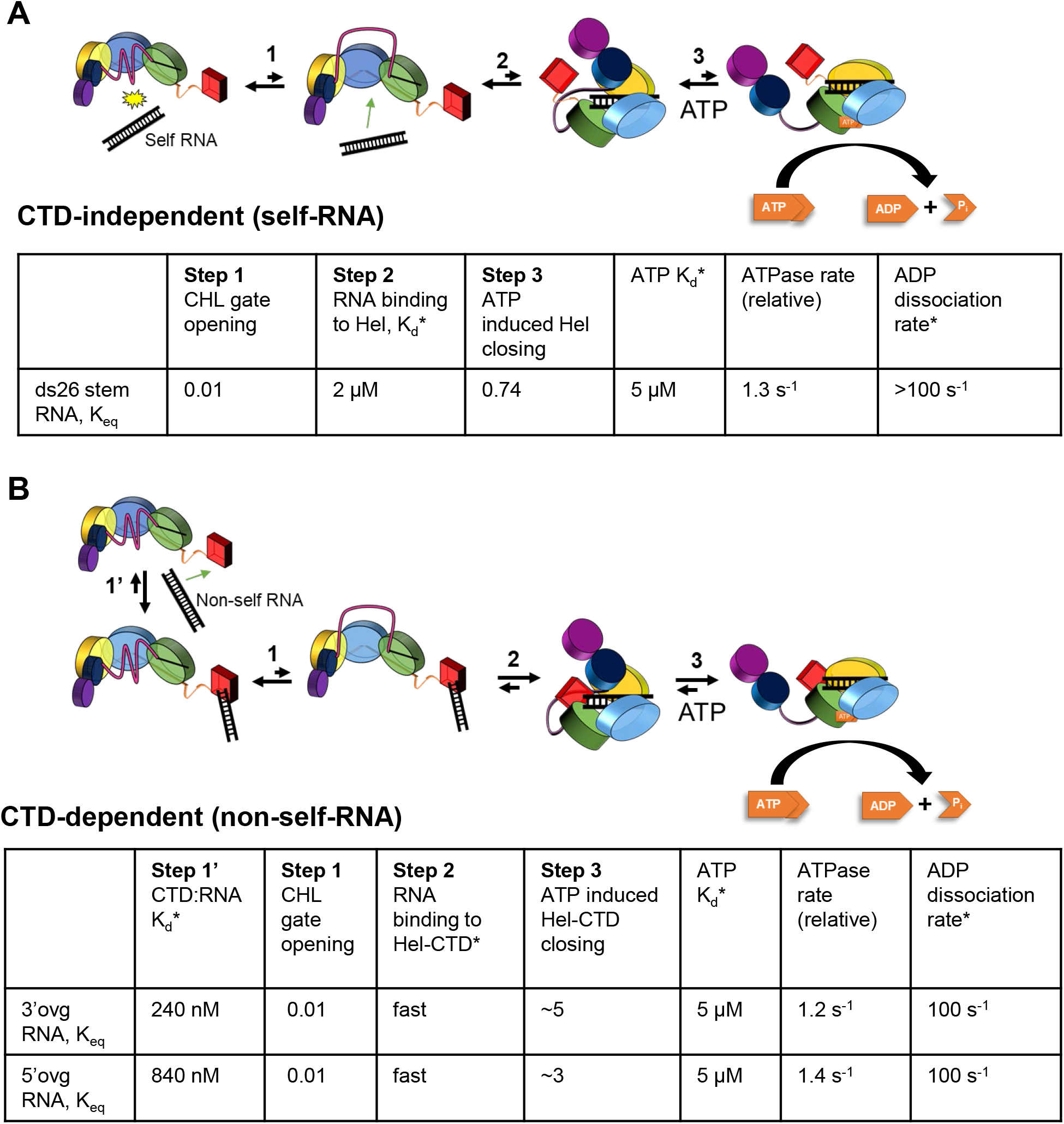
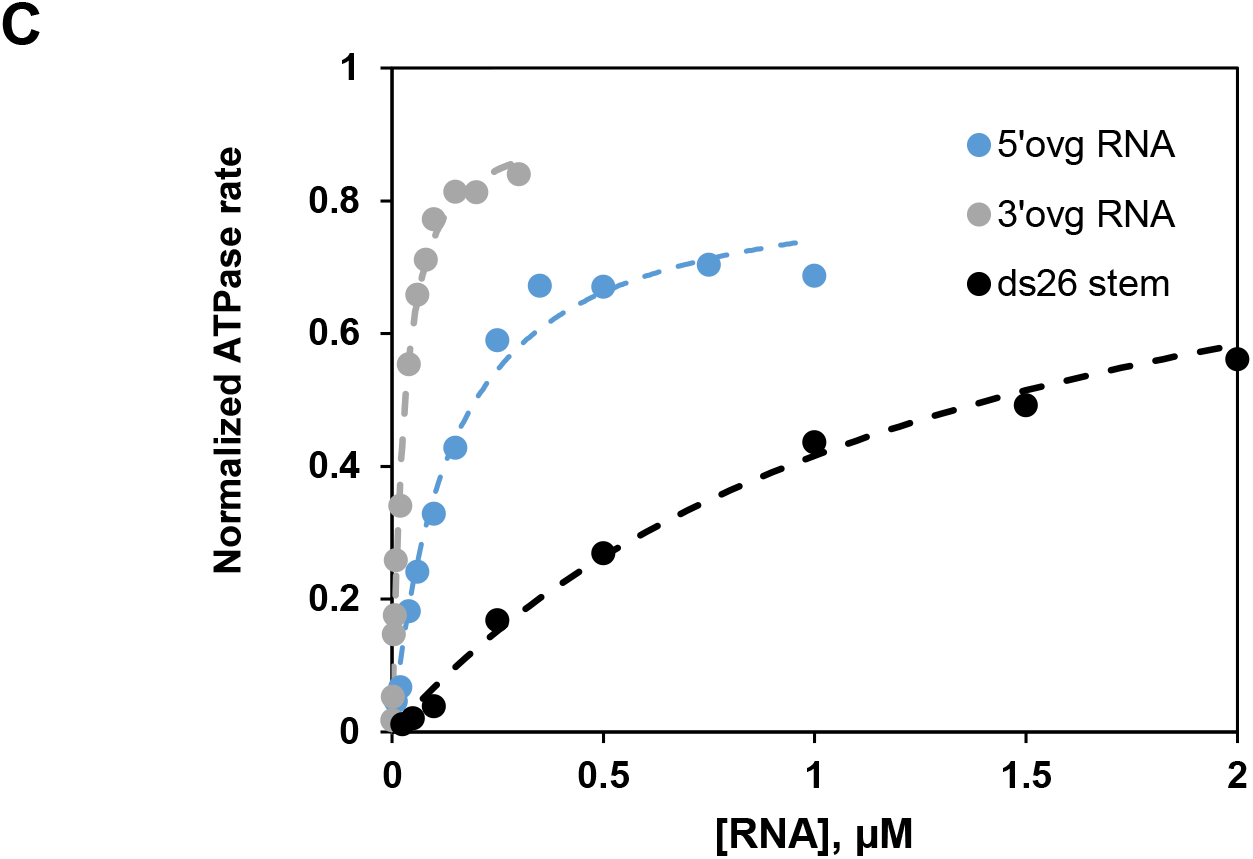
Kinetic modeling of RIG-I:RNA binding that includes the newly discovered role of the CHL in RIG-I regulation. (A) shows the model used to fit the ds26 stem RNA binding data of WT RIG-I using the KinTek Explorer software, and (B) shows the model to fit the 3’ovg and 5’ovg RNA binding data. In these models, the apo WT RIG-I exists in the autoinhibited state (with Hel2-Hel2i rotated and CARDs autoinhibited). In this state, the CHL-gate equilibrates between gate-open and gate-closed states. The CTD is free to bind the RNA in gate-closed autoinhibited RIG-I state, but the helicase domain binds RNA only in the CHL gate-open state. RNA binding induces conformational changes, first to a semiclosed helicase conformation and then to closed helicase conformation. The closed helicase conformation has exposed CARDs and is competent in ATP hydrolysis. The helicase can undergo multiple ATPase turnover as long as RNA remains bound. Best fit parameters of WT RIG-I binding to ds26 stem, 5’ovg RNA, and 3’ ovg RNA are shown in the Tables. The * denotes the parameters that were fixed to the experimental or the measured values during the fittings. (C) The RNA binding experimental data (circles from Figure 5) overlaid with the simulated curves (dashed lines) generated from the models and parameters in (A) and (B). Note, the K_eq_ values of step 1 and step 3 are dependent parameters (related to Figure 6).

**Table S1.**
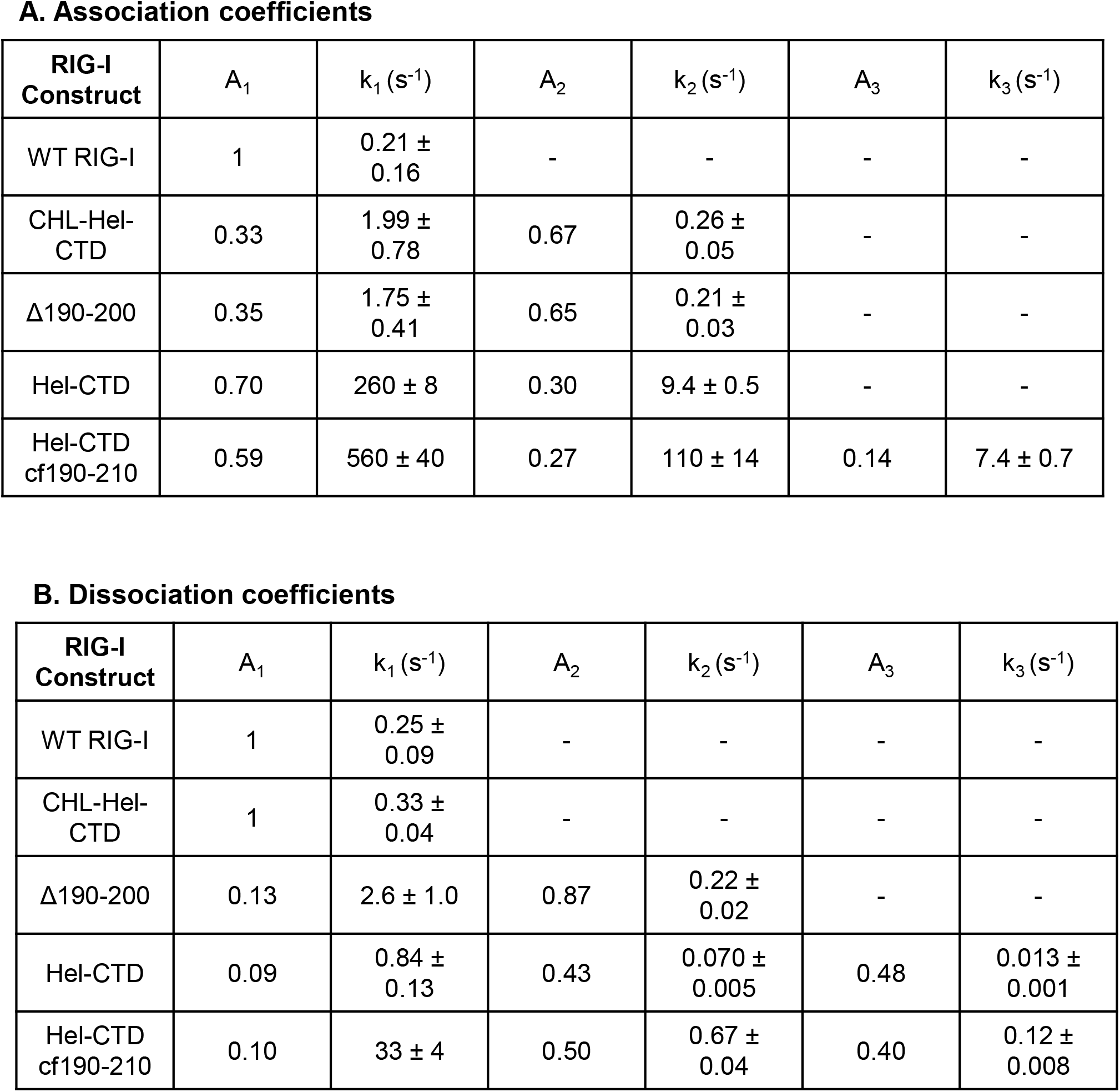
The stopped-flow kinetics of RNA association (A) and dissociation (B) experiments shown in Figure EV4 were fit to one or the sum of two or three exponentials and the associated coefficients are shown. A_n_ refers to the population fraction and k_n_ the rate constant. Each fit is an average of at least 6 individual traces. Standard errors of the fit are shown (related to Figure 4).

A. Association coefficients
| RIG-I Construct | $A_1$ | $k_1 (s^{-1})$ | $A_2$ | $k_2 (s^{-1})$ | $A_3$ | $k_3 (s^{-1})$ |
| --- | --- | --- | --- | --- | --- | --- |
| WT RIG-I | 1 | $0.21 \pm 0.16$ | - | - | - | - |
| CHL-Hel-CTD | 0.33 | $1.99 \pm 0.78$ | 0.67 | $0.26 \pm 0.05$ | - | - |
| $\Delta 190-200$ | 0.35 | $1.75 \pm 0.41$ | 0.65 | $0.21 \pm 0.03$ | - | - |
| Hel-CTD | 0.70 | $260 \pm 8$ | 0.30 | $9.4 \pm 0.5$ | - | - |
| Hel-CTD cf190-210 | 0.59 | $560 \pm 40$ | 0.27 | $110 \pm 14$ | 0.14 | $7.4 \pm 0.7$ |

**Table S1.** The stopped-flow kinetics of RNA association (A) and dissociation (B) experiments shown in Figure EV4 were fit to one or the sum of two or three exponentials and the associated coefficients are shown.
| RIG-I Construct | $A_1$ | $k_1 (s^{-1})$ | $A_2$ | $k_2 (s^{-1})$ | $A_3$ | $k_3 (s^{-1})$ |
| --- | --- | --- | --- | --- | --- | --- |
| WT RIG-I | 1 | $0.25 \pm 0.09$ | - | - | - | - |
| CHL-Hel-CTD | 1 | $0.33 \pm 0.04$ | - | - | - | - |
| $\Delta 190-200$ | 0.13 | $2.6 \pm 1.0$ | 0.87 | $0.22 \pm 0.02$ | - | - |
| Hel-CTD | 0.09 | $0.84 \pm 0.13$ | 0.43 | $0.070 \pm 0.005$ | 0.48 | $0.013 \pm 0.001$ |
| Hel-CTD cf190-210 | 0.10 | $33 \pm 4$ | 0.50 | $0.67 \pm 0.04$ | 0.40 | $0.12 \pm 0.008$ |

**Table S2.** Steady state RIG-I:RNA affinity constants. RNA K_D,app_ values reported as bars in Figure 3C-E. Standard errors are derived from the errors of fit for two binding trials (related to Figure 5).

| $K_{D, app}$ (nM) | 3'ovg RNA | 5'ovg RNA | stem RNA |
| --- | --- | --- | --- |
| **WT RIG-I** | 23.1 ± 5.5 | 105 ± 32 | 1250 ± 183 |
| **CHL-Hel-CTD** | 3.6 ± 2.9 | 4.9 ± 0.8 | 42.0 ± 13.6 |
| **Hel-CTD** | 0.27 ± 0.18 | 0.4 ± 0.4 | 2.0 ± 0.8 |
| **cf190-210 CHL-Hel-CTD** | 1.0 ± 1.4 | 0.80 ± 0.85 | 2.8 ± 2.1 |

**Table S3.** Energetic contributions of RIG-I domains to autoinhibition. Energetic contributions of CHL and CHL-CARDs, calculated through binding constants in Table S3. Equations used to calculate energetic contributions are Equations 7, 8, and 9 (related to Figure 5).

| Energetic contribution to autoinhibition ($\Delta\Delta G$) | 3'ovg RNA ($\Delta\Delta G$) kcal/mol | 5'ovg RNA ($\Delta\Delta G$) kcal/mol | stem RNA ($\Delta\Delta G$) kcal/mol |
| --- | --- | --- | --- |
| **CHL<sup>1</sup>** | -1.42 | -1.482 | -1.622 |
| **CHL-CARDs<sup>2</sup>** | -2.432 | -3.252 | -3.91 |

